# IFN-*γ* and TNF-*α* drive a *CXCL10*+ *CCL2*+ macrophage phenotype expanded in severe COVID-19 and other diseases with tissue inflammation

**DOI:** 10.1101/2020.08.05.238360

**Authors:** Fan Zhang, Joseph R. Mears, Lorien Shakib, Jessica I. Beynor, Sara Shanaj, Ilya Korsunsky, Aparna Nathan, Accelerating Medicines Partnership Rheumatoid Arthritis and Systemic Lupus Erythematosus (AMP RA/SLE) Consortium, Laura T. Donlin, Soumya Raychaudhuri

**Affiliations:** Center for Data Sciences, Brigham and Women’s Hospital, Boston, MA 02115, USA; Division of Genetics, Department of Medicine, Brigham and Women’s Hospital, Boston, MA 02115, USA; Department of Biomedical Informatics, Harvard Medical School, Boston, MA 02115 USA; Broad Institute of MIT and Harvard, Cambridge, MA 02142, USA; Division of Rheumatology, Inflammation, and Immunity, Brigham and Women’s Hospital and Harvard Medical School, MA 02115, USA; Graduate Program in Physiology, Biophysics and Systems Biology, Weill Cornell Graduate School of Medical Sciences, New York, NY 10065, USA; Arthritis and Tissue Degeneration, Hospital for Special Surgery, New York, NY, USA; Arthritis Research UK Centre for Genetics and Genomics, Centre for Musculoskeletal Research, The University of Manchester, Manchester, UK

## Abstract

Immunosuppressive and anti-cytokine treatment may have a protective effect for patients with COVID-19. Understanding the immune cell states shared between COVID-19 and other inflammatory diseases with established therapies may help nominate immunomodulatory therapies. Using an integrative strategy, we built a reference by meta-analyzing > 300,000 immune cells from COVID-19 and 5 inflammatory diseases including rheumatoid arthritis (RA), Crohn’s disease (CD), ulcerative colitis (UC), lupus, and interstitial lung disease. Our cross-disease analysis revealed that an *FCN1*+ inflammatory macrophage state is common to COVID-19 bronchoalveolar lavage samples, RA synovium, CD ileum, and UC colon. We also observed that a *CXCL10*+ *CCL2*+ inflammatory macrophage state is abundant in severe COVID-19, inflamed CD and RA, and expresses inflammatory genes such as *GBP1, STAT1*, and *IL1B*. We found that the *CXCL10*+ *CCL2*+ macrophages are transcriptionally similar to blood-derived macrophages stimulated with TNF-*α* and IFN-*γ ex vivo*. Our findings suggest that IFN-*γ*, alongside TNF-*α*, might be a key driver of this abundant inflammatory macrophage phenotype in severe COVID-19 and other inflammatory diseases, which may be targeted by existing immunomodulatory therapies.

## Introduction

Tissue inflammation is a unifying feature across diseases. While tissue- and disease-specific factors shape distinct inflammatory microenvironments, seemingly unrelated diseases can respond to the same therapy. For example, anti-tumor necrosis factor (TNF) therapies have revolutionized treatment for joint inflammation in autoimmune rheumatoid arthritis (RA) ^1^, while intestinal inflammation in Crohn’s Disease (CD) and ulcerative colitis (UC), collectively known as inflammatory bowel disease (IBD), also respond to anti-TNF medications ^2^. Here, we posit that deconstruction and subsequent integration of inflamed tissues at the level of individual cell phenotypes could provide a platform to identify shared pathologic features across diseases and provide rationale for repurposing medications in outwardly dissimilar conditions.

Recent studies have detailed features of local inflammation and immune dysfunction in COVID-19 and related diseases caused by SARS and MERS coronaviruses ^3^. Consensus is building that extensive unchecked inflammation involving so-called “cytokine storm” is a driver of severe late-stage disease. Single-cell studies of bronchoalveolar lavage fluid (BALF) have identified two inflammatory macrophage subsets characterized by expression of *FCN1* and *S100A8*, and *CCL2, CCL3*, and *CXCL10*, respectively, suggesting these cells might be high-level mediators of pathology in this late-stage disease ^4^. These macrophage subsets correlate with elevated circulating cytokine levels and extensive damage to the lung and vascular tissue. Independently, reports using peripheral blood from large numbers of COVID-19 patients have consistently documented lymphopenia (reduced lymphocyte frequency) paired with increased monocytes and inflammatory cytokines ^5–7^. Recent data suggest that moderate COVID-19 patients recovery associates with elevated tissue healing programs and lymphocyte growth factors, where severe patients maintain increased monocyte levels in blood and specific cytokines such as IFN-*α*, IFN-*γ*, and TNF, which appear ineffective in lowering viral load while possibly contributing to cytokine release syndrome (CRS) pathology ^7^. Together, these studies indicate the importance of uncovering the full extent of cell states present in COVID-19 patients including within affected tissues, in particular for monocytes and macrophages. Further, the extent to which these cell states are shared between COVID-19 and other inflammatory diseases and their disease association may further clarify disease mechanisms and precisely define therapeutic targets.

Macrophages are pervasive throughout the body and pivotal to tissue homeostasis, where they tailor their function to the parenchymal needs of each tissue type. In inflammation, tissue-resident macrophages and infiltrating monocytes are activated not only by factors from the unique tissue microenvironment, but yet additional layer of complexity elicited by disease-associating factors such as deregulated homeostatic byproducts, tissue damage, shifts in gene expression due to genetic variants, various immune cellular and soluble infiltrates and in some cases pathogenic microorganisms. The unprecedented plasticity and robust reactivity of macrophages and monocytes generates a spectrum of phenotypes yet to be fully defined in human disease that mediate clearance of noxious elements but in some cases, such as in cytokine storms, aggravates disease pathology. These include a range of pro-inflammatory and anti-microbial states that secrete key cytokines (e.g. TNF and IL-1 *β*) and chemokines (e.g. CXCL10 and CXCL11) and other functional states geared towards debris clearance, dampening inflammation and tissue reconstruction with factors such as MERTK, IL-10 and TGF*β*, respectively, as well as a variety of intermediate states ^8–10^. However, the full extent of shared immune cell states and secreted cytokines and chemokines, especially within activated macrophages that fuel inflammation, are so far unclear. Meta-analysis of the reactive macrophage phenotypes in inflamed tissues across diseases may further refine our understanding of the complexity of human macrophage function, while identifying inflammatory macrophage subsets potentially shared across multiple immune disorders and COVID-19, therein potentially providing a direct route to promising repurposed therapeutic strategies.

Single-cell RNA-seq (scRNA-seq) has provided an opportunity to interrogate inflamed tissues and identify pathogenic immune cell types ^11^. We recently defined a distinct *CD14*+ *IL1B*+ pro-inflammatory macrophage population that is markedly expanded in RA compared to osteoarthritis (OA), a non-inflammatory disease ^12,13^. Likewise, scRNA-seq studies on inflamed colonic tissues have identified inflammatory macrophage and fibroblast phenotypes with high levels of OSM signaling factors that are associated with resistance to anti-TNF therapies ^14^. Only very recently, developments in computational methods have made it possible to meta-analyze an expansive number of cells across various tissue states, while mitigating experimental and cohort-specific artifacts ^15–21^, therein assess shared and distinct cell states in disparately inflamed tissues.

To define the key shared immune cell compartments between inflammatory diseases with COVID-19, we meta-analyzed and integrated tissue-level single-cell profiles from 6 inflammatory diseases and COVID-19. We created an immune cell reference consisting of 307,084 single-cell profiles from 125 donor samples from RA synovium, systemic lupus erythematosus (SLE) kidney, UC colon, CD ileum, interstitial lung disease, and COVID-19 BALF. This single-cell reference represents comprehensive immune cell types from different disease tissues with different inflammation levels, which can be used to investigate other inflammatory diseases and their connections with COVID-19 in terms of immune cell responses. Using our meta-dataset reference, we identified major immune cell lineages including macrophages and monocytes, dendritic cells, T cells, B cells, NK cells, plasma cells, mast cells, and cycling lymphocytes. Among these, we found two inflammatory *CXCL10*+ *CCL2*+ and *FCN1*+ macrophage states that are shared between COVID-19 and inflammatory diseases. To understand the factors driving these phenotypes, we stimulated human blood-derived macrophages with eight different combinations of inflammatory disease-associated cytokines and tissue-associating stromal cells and analyzed it by scRNA-seq. We demonstrated that the *CXCL10*+ *CCL2*+ macrophages from severe COVID-19 lungs share a transcriptional phenotype with macrophages stimulated by TNF-*α* plus IFN-*γ*. Most notably the other two conditions wherein these macrophages are most abundant are RA and CD. This potentially provides a proof-of-concept regarding the power in identifying shared cellular states in unrelated inflamed tissues that align with sensitivity to the same medication—as both RA and CD respond to anti-TNF therapies. Furthermore, janus kinase (JAK) inhibitors have also proved effective in RA, in likely by targeting IFN-*γ* responses, which may indicate late-stage cytokine storm COVID-19 disease may involve Type II interferon and TNF responses and blocking these responses in macrophages may be a plausible treatment approach.

## Results

### A reference of > 300,000 immune single-cell profiles across common inflammatory diseases and COVID-19

To compare hematopoietic cells across inflammatory diseases and COVID-19 in an unbiased fashion, we aggregated 307,084 single-cell RNA-seq profiles from 125 healthy or inflammatory disease-affected donors spanning 6 disorders: (1) colon biopsies from healthy individuals, UC patients with inflamed or non-inflamed tissues ^14^; (2) terminal ileum tissue from patients with inflamed or non-inflamed CD ^22^; (3) synovial tissue from patients with RA or OA ^12,23^; (4) kidney biopsies from patients with SLE or healthy controls ^24^, (5) lung tissue from patients with interstitial lung disease ^25^ and (6) BALF from healthy individuals, mild or severe COVID-19 infection ^4^ (**Figures 1a-b, Supplementary Figure 1a, Supplementary Table 1**). We developed a pipeline for multi-tissue integration at the single-cell data (**Figure 1a**, **Methods**). First, we obtained and aggregated raw count matrices into a uniform matrix, and performed consistent quality control (QC) and normalization and scaling (**Methods**). To account for different cell numbers from different datasets, we performed weighted principal component analysis (PCA) by assigning higher weights to the cells from the dataset with a relatively small number of cells and vice versa. Then, we used our batch-correction algorithm Harmony ^15^ to integrate these diverse datasets, accounting for variation due to different levels of technical and biological effects that confound cell type identification (**Methods**). To quantify the integration of the datasets, we employed the local inverse Simpson’s Index (LISI) ^21^. A LISI score of 1.0 means that there is no mixing, and higher scores indicate better mixing of donors and tissue sources (**Methods**). We observed that applying Harmony increased mixing among donors (LISI increasing from mean 2.9 to 6.1) and tissue sources (LISI increasing from mean 1.0 to 1.8, **Supplementary Figure 2a**).

**Figure 1.**
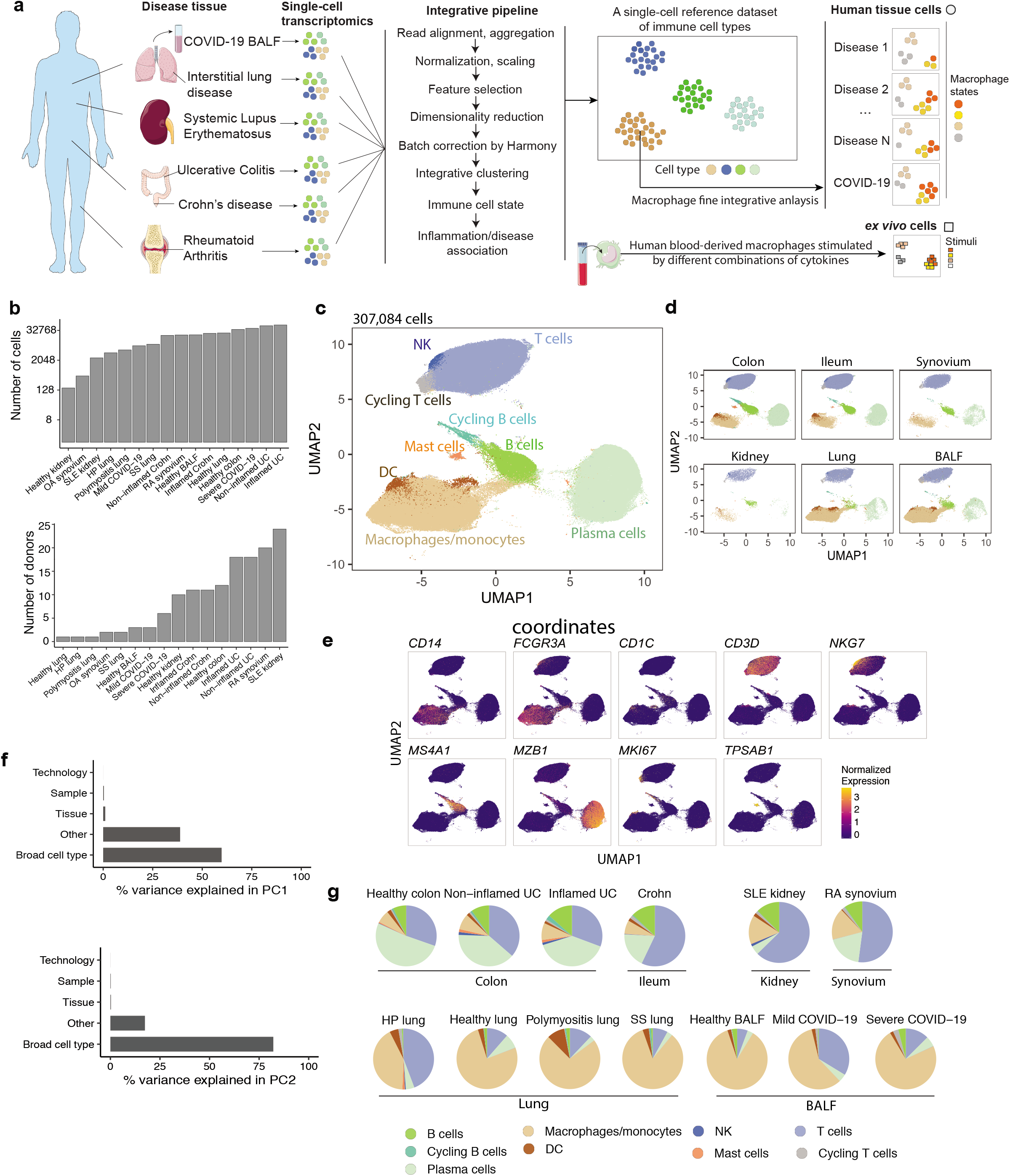
Integrative transcriptomic analysis of >300,000 single-cell profiles from 6 inflammatory disease tissues and COVID-19 reveals shared immune cell populations. **a**. Overall study design and single-cell analysis, including the integrative pipeline, a single-cell reference dataset, and *ex vivo* stimulated macrophage dataset. Shared states, specifically macrophages, are identified across disease tissues, and then compared to the *ex vivo* cells to identify the stimuli driving their phenotype. **b**. Number of cells and donor samples from each healthy and disease tissue. SS lung denotes systemic sclerosis lung; HP lung denotes hypersensitivity pneumonitis lung. **c**. Integrative clustering of 307,084 cells reveals common immune cell types from different tissue sources. Cells from the same cell types are projected together in UMAP space. **d**. Immune cells from separate tissue sources in the same UMAP coordinates as in **c**. **e**. Expression of cell type lineage marker genes in the UMAP space. **f**. Percent of variance explained in the gene expression data by pre-defined broad cell types, donor samples, tissue sources, and technologies for the first and second principal component (PC1 and PC2) after batch effect correction. **g**. Proportions of identified immune cell types within each disease tissue or healthy control.

This approach enabled broad cell type categorization in the hematopoietic cell lineage. We performed graph-based clustering ^26^ on the integrated principal components (PCs) and dimensionality reduction using UMAP (Uniform Manifold Approximation and Projection) to project cells into two-dimensional space ^27^. We identified T cells (marked by *CD3D* expression), NK cells (*NKG7*), B cells (*MS4A1*), plasma cells (*MZB1*), macrophages (*FCGR3A*) and monocytes (*CD14*), dendritic cells (DCs)(*CD1C*), mast cells (*TPSAB1*), and cycling lymphocytes (*MKI67*) (**Figure 1c-e, Supplementary Figure 1b**).

Our cross-tissue integration pipeline successfully identified previously reported disease-specific patterns. This included the presence of germinal center B cells in the inflamed UC colon and age-associated B cells in RA synovium (**Supplementary Figure 1c**). Furthermore, we observed that the majority of variance (>60% in PC1 and PC2) derived from gene expression patterns are explained by major cell types (**Figure 1f, Supplementary Figure 1d**). In contrast, variables such as tissue type, technology, or donor sample accounted for <1% of the variation in PC1 and PC2 after batch effect correction. We note that prior to Harmony batch effect correction, the same cell types from different tissues fail to integrate together (**Supplementary Figure 2b**).

The integration of single-cell data across tissues from multiple diseases provided a means to quantify the contribution of distinct immune cell types to the various inflammatory conditions (**Figure 1g**). Proportions of major immune cell types residing in different tissue sources are different. Overall, samples obtained from lung tissues, whether from healthy controls or patients with different conditions, contained the highest proportion of macrophages (~73.2% of total hematopoietic cells). The RA synovium, SLE kidney, and CD ileum were dominated by T lymphocytes (57.3%), while the UC colon samples had a distinctively high abundance of plasma cells (43.3%) (**Figure 1g**). Severe COVID-19 bronchoalveolar lavage samples in comparison to mild COVID-19 also contained a higher proportion of macrophages similar to other lung tissues (**Figure 1g**). The large number of cells across multiple disease and tissue contexts positioned us to precisely characterize cell states (**Figure 1b-c**).

### Identification of shared inflammatory macrophage states across inflammatory disease tissues and COVID-19

Macrophages represented a dominant cell type across all affected target tissues ^12,14,22–25^. Therefore, we performed a fine clustering analysis on these cells to define shared and distinct macrophage states and phenotypes across these inflammatory diseases and COVID-19 (**Figure 2a**). To this end, we applied the same integrative pipeline on 74,373 macrophages and monocytes from synovium, ileum, colon, lung, and BALF from 108 individuals (**Supplementary Table 2**). We identified a total of four states: *CXCL10*+ *CCL2*+ *CD14*+ *FCGR3A*+ inflammatory macrophages, *FCN1*+ *CD14*+ *FCGR3A*+ inflammatory macrophages, M2-like anti-inflammatory *MRC1*+ *FABP4*+ macrophages, and non-inflammatory macrophages (**Figure 2a-b, Supplementary Figure 3a**). The two inflammatory macrophage states correspond to the previously identified *CXCL10*+ and *FCN1*+ macrophages in COVID-19 BALF, respectively ^4^. Notably, in this clustering, previously described inflammatory macrophages identified in inflamed RA synovium and in inflamed UC and CD intestinal tissue clustered along with the majority of the severe COVID-19 macrophages, which spanned across these two inflammatory *CXCL10*+ *CCL2*+ and *FCN1*+ states (**Figure 2c**, **Supplementary Figure 3b-c**). The LISI score that evaluates dataset mixing decreased with respect to previously described macrophage annotations, and increased with respect to donor- and tissue-specific effects after batch correction (**Supplementary Figure 3d**), indicating that the shared macrophage subsets were driven primarily by macrophage biology-related gene expression patterns rather than tissue or donor source.

**Figure 2.**
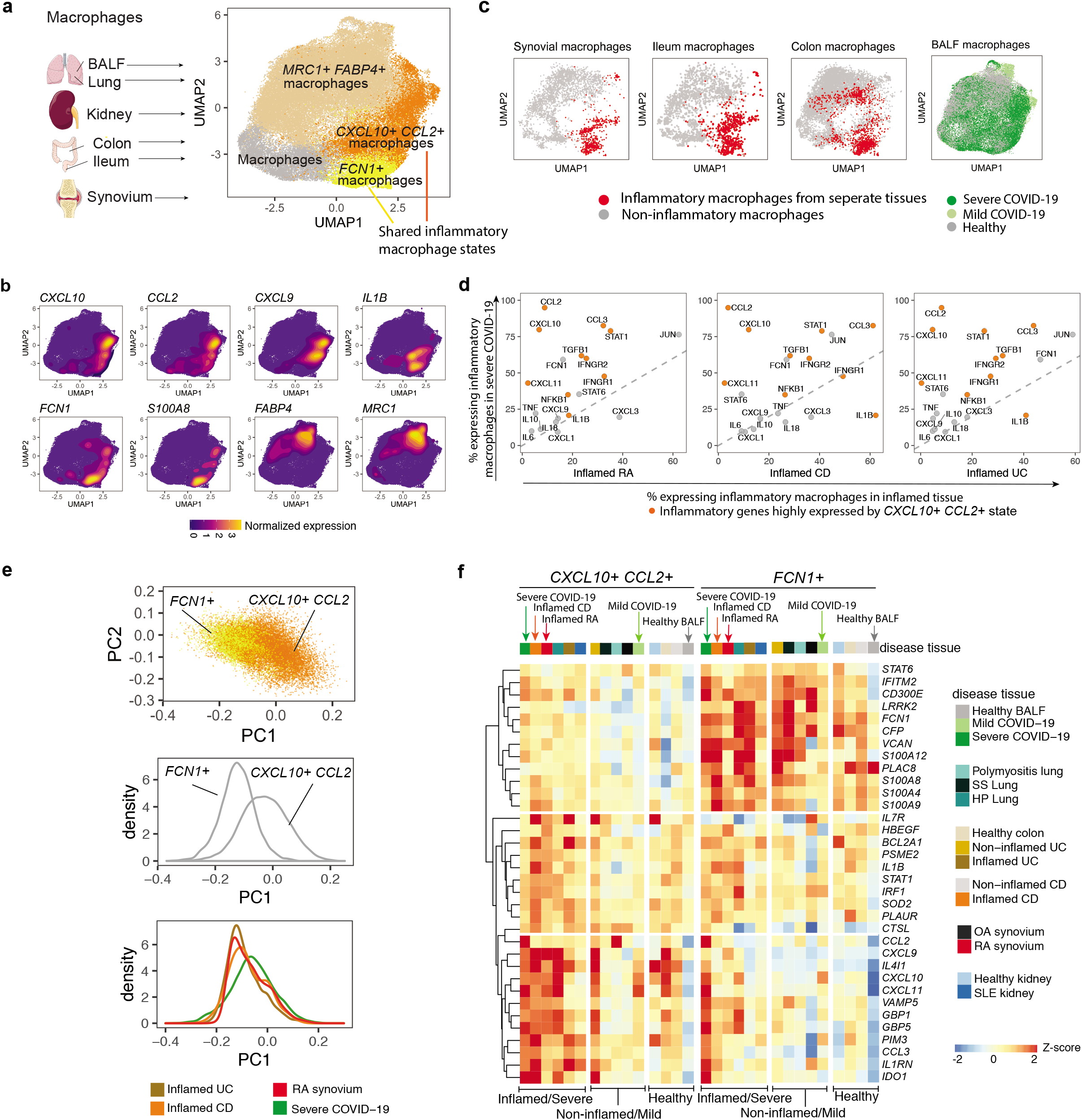
Integrative analysis of macrophages reveals shared *CXCL10*+ *CCL2*+ and *FCN1*+ inflammatory macrophage states across inflammatory disease tissues and COVID-19. **a**. Integrative clustering of 74,373 macrophages and monocytes from 108 individuals from BALF, lung, kidney, colon, ileum, and synovium reveals four distinct macrophage states. Two inflammatory macrophage states are observed: *CXCL10*+ *CCL2*+ and *FCN1*+ inflammatory macrophages. **b**. Density plot of cells with non-zero expression of cluster marker genes in UMAP space. **c**. Previously defined inflammatory macrophages from different inflammatory disease tissues are clustered together with the majority of the macrophages from severe COVID-19 in the integrative embeddings. Inflammatory macrophages are separated into the *CXCL10*+ *CCL2*+ and *FCN1*+ inflammatory states. **d**. Proportion of expressing (non-zero) inflammatory cytokines and genes from inflammatory macrophages in inflamed RA, CD, and UC compared to those in severe COVID-19. Genes that are highly expressed in the *CXCL10*+ *CCL2*+ inflammatory macrophages are highlighted in orange. **e**. PCA analysis on the identified inflammatory macrophages. The first PC captures a gradient from the *FCN1*+ state to the *CXCL10*+ *CCL2*+ state. Two distributions are shown to represent the density of the macrophages mapping to PC1. Macrophages from inflamed tissues are mapped to PC1 coordinates. A shift on PC1 loadings between inflammatory macrophages from inflamed UC and severe COVID-19 (Wilcoxon rank-sum test *P* < 2.2e-16), inflamed RA and severe COVID-19 (*P* = 0.001), and inflamed CD and severe COVID-19 (*P* = 1.4e-07) are displayed, respectively. **f**. Heatmap of Z-score of the average expression of top marker genes for the *CXCL10*+ *CCL2*+ and *FCN1*+ inflammatory macrophage states. Rows include genes and columns show pseudo-bulk expression per condition within each state. Gene signatures were selected based on AUC > 0.6 and Bonferroni-adjusted *P* < 10^−5^ comparing cells from one cluster to the others using pseudo-bulk analysis.

To further explore how the *CXCL10*+ *CCL2*+ and *FCN1*+ macrophages are involved in tissue inflammation, we examined key inflammatory features ^14^ that were expressed in these two states. A high proportion of inflammatory macrophages in severe COVID-19, RA, UC, and CD expressed inflammation-associated factors including *CXCL10, CXCL11, CCL2, CCL3, STAT1, IFNGR1, IFNGR2, NFKB1, TGFB1*, and *IL1B* (**Figure 2d, Supplementary Figure 4a**). The gene signature for the *CXCL10*+ *CCL2*+ inflammatory macrophage state was found in a higher proportion of macrophages in severe COVID-19 than in the other inflamed tissues (**Figure 2d**). Applying PCA to the two inflammatory macrophage states, we found that PC1 captured a gradient from the *FCN1*+ state to the *CXCL10*+ *CCL2*+ state (**Figure 2e**), suggesting a potential continuum with intermediates between the inflammatory *FCN1*+ and *CXCL10*+ *CCL2*+ states. While the majority of inflammatory macrophages in RA, UC, and CD align more closely with the *FCN1*+ state, in severe COVID-19 we observed a shift in cell frequency, corresponding to a higher abundance of *CXCL10*+ *CCL2*+ macrophages compared to other inflammatory diseases (**Figure 2e, Supplementary Figure 4b**).

To more extensively define markers for the two inflammatory tissue macrophage states shared across COVID-19, RA, UC and CD, we performed pseudo-bulk differential expression analysis (**Methods, Supplementary Table 3**, AUC > 0.6, Bonferroni-adjusted *P* < 10^−5^). The *CXCL10*+ *CCL2*+ inflammatory macrophages displayed significantly higher expression of *CXCL10, CXCL11, CCL2, CCL3, GBP1*, and *IDO1* in severe COVID-19, inflamed RA, and CD compared to *FCN1*+ macrophages (**Figure 2f**). The *FCN1*+ macrophages show high expression of *FCN1* (Ficolin-1) and alarmins *S100A8* and *S100A9* in most of the inflamed tissues compared to *CXCL10*+ *CCL2*+ inflammatory macrophages (**Figure 2f**). Both inflammatory macrophage states show high expression of M1 macrophage-related transcription factors, *STAT1* and *IRF1*, in inflamed RA, UC, CD, and COVID-19 BALF relative to healthy or non-inflamed tissues (**Figure 2f**). Within the *CXCL10*+ *CCL2*+ state, we noted heterogeneity that correlates with *IL1B* expression (**Supplementary Figure 4c-d**). Moreover, when we examined the effect size of all genes by comparing *CXCL10*+ *CCL2*+ and *FCN1*+ macrophages with *MRC1+ FABP4*+ macrophages within each tissue, inflammatory genes indeed demonstrated the greatest fold change differences (**Supplementary Figure 5**). We further examined these inflammation-associated genes using CD45+ CD14+ flow sorted bulk RNA-seq samples from inflamed (leukocyte-rich) RA, non-inflamed (leukocyte-poor) RA, and OA ^12^; we see the *CXCL10*+ *CCL2*+ state-specific genes (*CXCL10, CXCL9, CCL3, GBP1*, and *IDO1), FCN1*+ state-specific genes (*FCN1, S100A9, CD300E, IFITM3*, and *CFP*), and genes (*IRF1*, *BCL2A1*, and *STAT1*) associated with both states are significantly enriched in the macrophages from inflamed RA compared to non-inflamed RA and OA (**Supplementary Figure 6**). By integrating macrophages across multiple inflamed tissues, we show that inflammatory subsets identified in COVID-19 may share common phenotypes with macrophages from other inflammatory conditions.

### Tissue inflammatory conditions drive distinct macrophage phenotypes

To define the factors within tissues that collectively shape disease-associated macrophage states, we generated human blood-derived macrophages and activated them with eight mixtures of inflammatory factors, with particular interest in antiviral interferons (IFN-# and IFN-*γ*) and pro-inflammatory cytokines such as TNF that mediate CSR and mediates RA and IBD pathology (**Figure 3a, Supplementary Figure 7a-c, Methods**). We added fibroblasts in some conditions to mimic exposure to the stromal factors found within tissues. To experimentally minimize confounding batch effects during scRNA-seq barcode labeling, we used a single-cell antibody-based hashing strategy ^28^ to multiplex samples from different stimulatory conditions in one sequencing run. We used 9 hashtag antibodies on 4 donor samples (**Supplementary Table 4, 5**), and obtained 25,823 post-QC cells after applying to 10X Genomics droplet-based single-cell assay (**Supplementary Figure 7b-c, Methods**).

**Figure 3.**
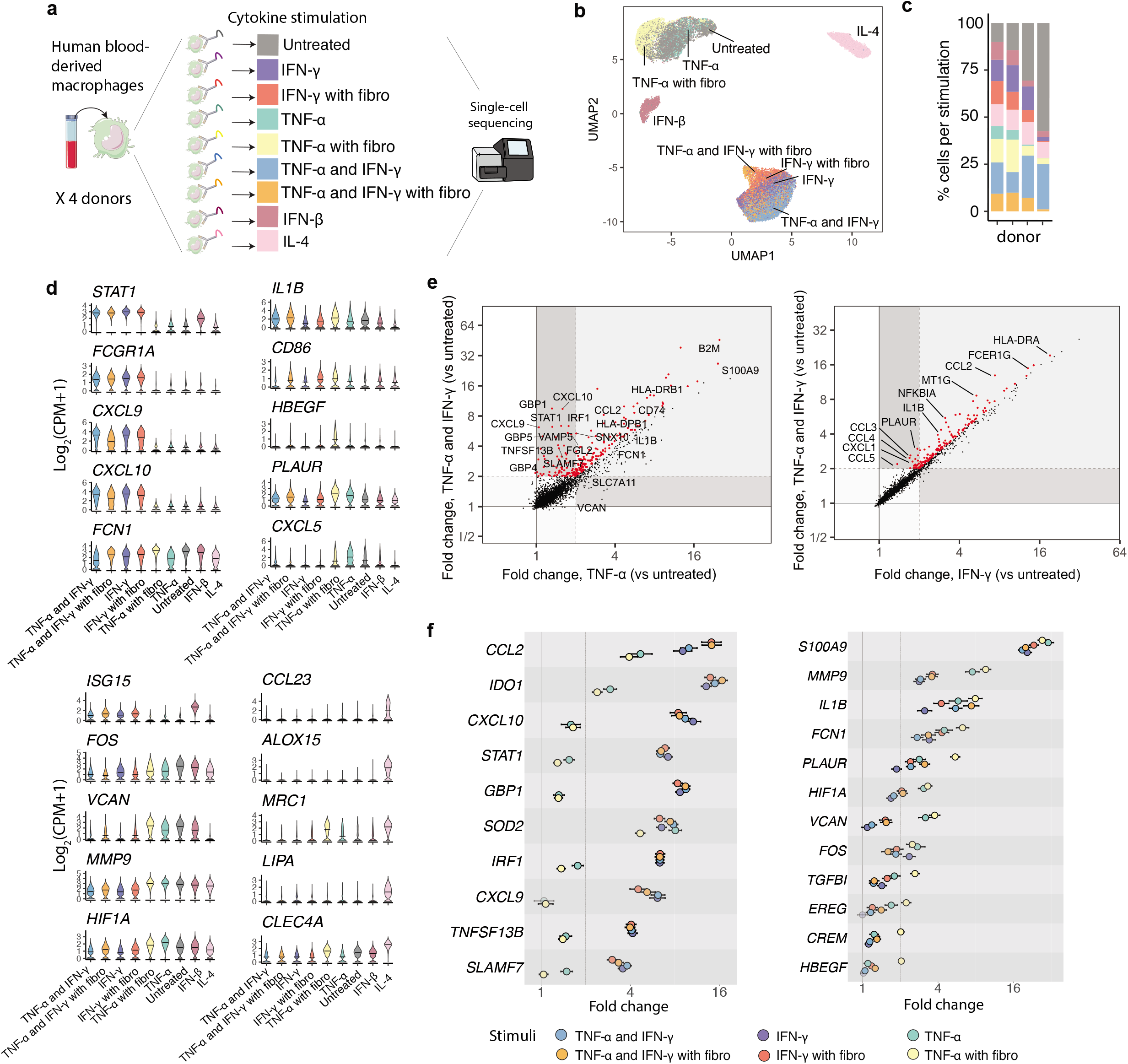
Human blood-derived macrophages stimulated by eight mixtures of inflammatory factors present heterogeneous macrophage phenotypes. **a**. Schematic representation of the single-cell cell hashing experiment on human blood-derived macrophages stimulated by eight mixtures of inflammatory factors from 4 donor samples. A diagram of the single-cell antibody-based hashing strategy used to multiplex samples from different stimulatory conditions in one sequencing run. Here fibro denotes fibroblasts. **b**. Condition labels of the stimulated 25,823 blood-derived macrophages from 4 donor samples are colored and labeled in UMAP space. **c**. Proportion of different stimulatory conditions for each donor sample are calculated. **d**. Log-normalized expressions of genes that are specific to different conditions are displayed in violin plots. Mean of normalized gene expression is marked by a line and each condition by individual coloring. CPM denotes counts per million. **e**. Fold changes in gene expression after TNF-*α* stimulation vs. TNF-*α* and IFN-*γ* stimulation (left), and IFN-*γ* vs. TNF-*α* and IFN-*γ* stimulation (right) for each gene. Genes in red have fold change > 2, Bonferroni-adjusted *P* <10^−7^, and a ratio of TNF-*α* and IFN-*γ* fold change to TNF-*α* fold change greater than 1 (left) or a ratio of TNF-*α* and IFN-*γ* fold change to IFN-*γ* fold change greater than 1. Genes that are most responsive to either IFN-*γ* (left) or TNF-*α* (right) are labeled. **f**. Stimulation effect estimates of genes that are most responsive to conditions with IFN-*γ* or TNF-*α* with fibroblasts comparing each condition to untreated macrophages using linear modeling. Fold changes with 95% CI are shown.

We produced single-cell expression profiles for stimulated blood-derived macrophages and labeled them by their conditions (**Figure 3b-c**). Consistent with well-established effects, macrophages stimulated by IL-4 show increased expression of *CCL23, MRC1*, and *LIPA*, markers of the M2-like anti-inflammatory state (**Figure 3d**). Differential expression analysis revealed that all conditions containing IFN-*γ* (Type II Interferon) resulted in macrophages with high levels of the transcription factor *STAT1*, interferon-stimulated genes *CXCL9* and *CXCL10* and inflammatory receptors such as *FCGR1A ^29^* (**Figure 3d**).

Using linear models, we identified genes with the greatest response to each stimulation and estimated their effect sizes (**Methods**). We found 403 genes (Fold change > 2, FDR < 0.05) that were significantly enriched in the TNF-*α* and IFN-*γ* stimulation compared to untreated macrophages. Furthermore, a group of genes including *CCL2*, *CXCL9, CXCL10, SLAMF7*, and *STAT1* had a significantly higher induction in macrophages exposed to TNF-*α* and IFN-*γ* stimulation compared to TNF-*α* alone (**Figure 3e** left). We observed similar effect sizes for these genes when we stimulated macrophages with TNF-*α* and IFN-*γ*, TNF-*α* and IFN-*γ* with fibroblasts, IFN-*γ*, and IFN-*γ* with fibroblasts compared to untreated macrophages (**Figure 3f** left). Other stimulatory conditions with TNF-*α* only or TNF-*α* with fibroblasts show no or substantially less expression induction of these genes (**Figure 3f** left, **Supplementary Figure 7d**). We consider these genes to reflect a specific IFN-*γ*-driven signature. We also observed a modest induction of TNF-*α*-driven genes such as *CCL2*, *CCL3*, *IL1B*, and *NFKBIA* enriched in TNF-*α* and IFN-*γ* stimulation compared to IFN-*γ* alone (**Figure 3e** right). Additionally, we identified 400 genes (Fold change > 2, FDR < 0.05) including inflammatory regulators such as *FCN1* and *PLAUR* that are most highly induced in response to TNF-*α* stimulation with fibroblasts compared to no treatment (**Figure 3f** right). Overall, these findings indicated that TNF-*α*-driven responses appeared more malleable when combined with other factors, for example, wherein co-cultured fibroblasts enhanced TNF-*α*-induced *MMP9, PLAUR*, and *TGFBI*, while IFN-*γ* repressed this TNF-*α* effect. Notably, TNF-*α* and IFN-*γ* generated a macrophage phenotype with preserved expression of NF-kB targets such as *IL1B, NFKBIA*, and *HLA-DRA* together with STAT1 targets such as *CXCL9* and *CXCL10*, and *GBP1* and *GBP5* (**Figure 3e-f**).

### Identification of a TNF-*α* and IFN-*γ* synergistically driven inflammatory macrophage phenotype expanded in severe COVID-19 and other inflamed disease tissues

Our cross-tissue integrative analysis revealed two shared inflammatory macrophage states (**Figure 2**). To further understand these cell states and the *in vivo* inflammatory tissue factors driving them, we integrated the single-cell transcriptomes of the tissue macrophages with the experimental multifactor-stimulated macrophages. After correcting for tissue source and donor effects, we identified 7 distinct macrophage clusters (**Figure 4a**). The tissue *CXCL10*+ *CCL2*+ inflammatory macrophages from UC colon, CD ileum, RA synovium, and COVID-19 BALF were transcriptionally most similar to macrophages stimulated by the combination of TNF-*α* plus IFN-*γ* in cluster 1 (**Figure 4b-c, Supplementary Figure 8a-c**). The blood-derived macrophages in cluster 1 include macrophages stimulated by four different conditions all including IFN-*γ*, of which 37.5% are macrophages stimulated by TNF-*α* and IFN-*γ* together (**Figure 4c, d**, **Supplementary Figure 8b**).

**Figure 4.**
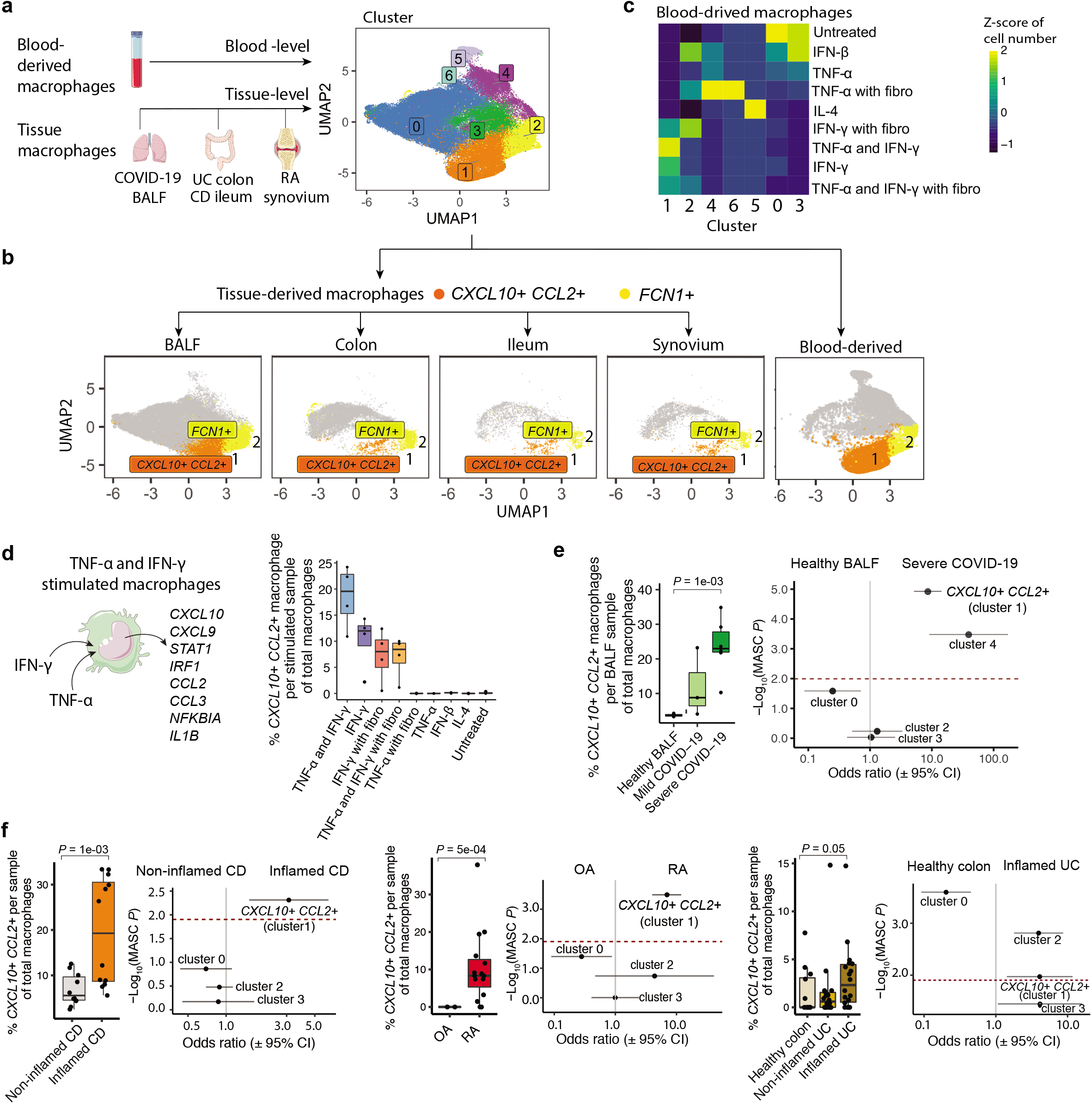
Identification of TNF-*α* and IFN-*γ* driven CXCL10+ CCL2+ inflammatory macrophages expanded in severe COVID-19 and other inflamed disease tissues. **a**. Integrative clustering of stimulated blood-derived macrophages with tissue-level macrophages from COVID-19 BALF, UC colon, CD ileum, and RA synovium. **b**. The previously identified tissue-level *CXCL10*+ *CCL2*+ state corresponds to cluster 1 (orange), and the *FCN1*+ inflammatory macrophage state corresponds to cluster 2 (yellow). Macrophages from each tissue source are displayed separately in the same UMAP coordinates as in **a**. **c**. Heatmap indicates the concordance between stimulatory conditions and cluster assignments. Z-score of the number of cells from one stimulatory condition to each of the clusters is shown. **d**. For the blood-derived stimulated macrophages, the proportion of *CXCL10*+ *CCL2*+ macrophages per stimulated donor sample of total macrophages are shown. **e** and **f**. For each tissue source, we show the proportion of *CXCL10*+ *CCL2*+ macrophages per sample of total macrophages from healthy BALF (*n* = 3), mild (*n* = 3) and severe (*n* = 6) COVID-19 BALF, non-inflamed CD (*n* = 10) and inflamed CD (*n* = 12), OA (*n* = 2) and RA (*n* = 15), and healthy colon (*n* = 12), non-inflamed (*n* = 18) and inflamed UC (*n* = 18). Medians of proportions for each group are shown. *P* is calculated by Wilcoxon rank-sum test within each tissue source. For each tissue source, the association of each cluster with severe/inflamed compared to healthy control was tested. 95% CI for the odds ratio (OR) is given for each cluster. MASC *P* is calculated based on one-sided F tests conducted on nested models with MASC ^30^. The clusters above the dashed red line (MASC *P* threshold after Bonferroni correction) are statistically significantly associated with inflammation/severity compared to non-inflammatory/healthy status. Clusters that have less than 30 cells are removed from association testing.

To elucidate cell states that were phenotypically associated, we tested the association of each cluster with severe COVID-19 compared to healthy BALF using a logistic regression model accounting for technical cell-level and donor-level effects ^30^ (**Methods**). We observed two clusters abundant in severe COVID-19 compared to healthy BALF: *CXCL10*+ *CCL2*+ (cluster 1), which is transcriptionally similar to the TNF-*α* and IFN-*γ* induced phenotype and cluster 4, which most closely matches a TNF-*α* with fibroblasts induced phenotype (**Figure 4e**). The *CXCL10*+ *CCL2*+ inflammatory macrophages are significantly more abundant in severe COVID19 (23.7%) compared to healthy BALF (3.7%), and express high levels of the genes that synergistically respond to TNF-*α* and IFN-*γ* stimulation (**Figure 4d-e, Supplementary Figure 8d-e**). We examined other diseases also, and observed that the *CXCL10*+ *CCL2*+ inflammatory macrophages are expanded in inflamed CD compared to non-inflamed CD, RA compared to non-inflammatory OA, and inflamed UC compared to healthy colon, respectively (**Figure 4f**). Taken together, these results indicate that TNF-*α* and IFN-*γ* drive the synergistic inflammatory response in the *CXCL10*+ *CCL2*+ inflammatory macrophage phenotype that is expanded not only in COVID-19, but also in inflamed tissues from other diseases, which suggests this inflammation-associated macrophage state may present a viable target for these diseases.

## Discussion

Our study demonstrates the power of a multi-disease reference dataset to interpret cellular phenotypes and tissue states, while placing them into a broader context that may provide insights into disease etiology and rationale for repurposing medications. Such meta-datasets can increase the resolution of cell states and abet understanding of shared cellular states found in less well-understood diseases such as COVID-19. Amassing diverse tissues from > 120 donors with a wide range of diseases, we built a human tissue inflammation single-cell reference. Applying powerful computational strategies, we integrate > 300,000 single-cell transcriptomes and correct for factors that interfere with resolving cell-intrinsic expression patterns. In particular, we have identified a *CXCL10*+ *CCL2*+ inflammatory macrophage phenotype shared between tissues affected in autoimmune disease (RA), inflammatory diseases (CD and UC), and infectious disease (COVID-19). We observed that the abundance of this population is associated with inflammation and disease severity. With integrated analysis of an *ex vivo* dataset, we elucidated its potential cytokine drivers: TNF-*α* together with IFN-*γ*.

Macrophages are ideal biologic indicators for the *in vivo* state of a tissue due to their dynamic nature, robust responses to local factors and widespread presence in most tissues. Through our cross-disease analysis, we defined two inflammatory macrophage states that can be found in selected groups of seemingly unrelated tissues and diseases. Most notably, the *CXCL10*+ *CCL2*+ inflammatory macrophages predominate in the bronchoalveolar lavage of patients with severe COVID-19, and are also seen in synovial tissue affected by RA and inflamed intestines. These cells are distinguished by high levels of *CXCL10* and *CXCL11, STAT1, IFNGR1* and *IFNGR2*, as well as, *CCL2* and *CCL3*, *NFKB1, TGFB1*, and *IL1B*. This gene expression pattern of the JAK/STAT and nuclear factor-κB (NF-kB) dependent cytokines implicates induction by an intriguing combination of both the IFN-induced JAK/STAT and TNF-induced NF-kB pathways and, in conjunction, the overall transcriptome program most closely aligns with macrophages stimulated by IFN-*γ* plus TNF-*α*. As both JAK inhibitors and anti-TNF medications have outstanding efficacy in treating RA and anti-TNFs are the most common medications treating inflammatory bowel disease including Crohn’s Disease ^2^, these therapies may target the inflammatory macrophages in severe COVID-19 lung during the phase involving a cytokine release syndrome ^31^.

Infection with SARS-CoV2 triggers local immune response and inflammation in the lung compartment, recruiting macrophages and monocytes that release and respond to inflammatory cytokines and chemokines ^6^. This response may change with disease progression, in particular during the transition towards cytokine storm associated with severe disease. Intriguingly, our cross-disease tissue study strongly suggests that IFN-*γ* is an essential component in the inflammatory macrophage phenotype in severe COVID-19. Most studies on the interferons and coronaviruses have focused on Type I Interferons, such as IFN-#, due to their robust capacity to interfere with viral replication ^32^. Indeed, ongoing research into the administration of recombinant IFN-# has shown promise in reducing the risk of severe COVID-19 disease ^33^. However, other studies have indicated that targeting IFN-*γ* may be an effective treatment for cytokine storm, a driver of severe disease in COVID-19 patients ^34,35^. Additionally, recent research has indicated that targeting IFN-*γ* using JAK inhibitors such as ruxolitinib, baricitinib, and tofacitinib offers effective therapeutic effects in treating severe COVID-19 patients ^31,36,37^. Clinical trials of Type II interferon inhibitors in COVID-19 are under way (NCT04337359, NCT04359290, and NCT04348695) ^31^. In agreement with these studies, our findings may indicate that IFNy is an important mediator of severe disease, in part through activating the inflammatory *CXCL10*+ *CCL2*+ macrophage subset. We hypothesize that anti-Type II interferon treatment, including JAK inhibitors, might prove effective at inhibiting the cytokine storm driving acute respiratory distress syndrome in patients with severe COVID-19. Of course, the presence of an IFN-*γ* and TNF-*α* phenotype is an association may not be causal. Whether targeting these cytokines is reasonable or not, will depend on additional clinical investigation.

## Acknowledgements

This work is supported in part by funding from the National Institutes of Health (NIH) Grants UH2AR067677, U01 HG009379, and 1R01AR063759 (to S.R.); and NIH R01AI148435, UH2 AR067691, Carson Family Trust, and Leon Lowenstein Foundation (to L.T.D.). We thank the BWH Single Cell Genomics Core for assistance in the single-cell hashing experiment. We thank members of the Raychaudhuri laboratory for discussions.

## Author contributions

S.R. and F.Z. conceptualized the study. F.Z. and S. R. designed the statistical and computational strategy, and analyzed the data. J.R.M. collected public single-cell datasets and conducted additional analyses. F.Z., S.R., and J.R.M. wrote the initial manuscript. L.T.D., A.N., I.K., J.I.B., L.S., and S.S. edited the draft. L.T.D obtained blood samples from human subjects. L.T.D, L.S., J.I.B., and S.S. organized processing, transportation, and experiment of the blood samples. S.R. and L.T.D. supervised the work. All authors discussed the results and commented on the manuscript.

## Accelerating Medicines Partnership Rheumatoid Arthritis & Systemic Lupus Erythematosus (AMP RA/SLE) Consortium

Jennifer Albrecht^9^, Jennifer H. Anolik^9^, William Apruzzese^5^, Brendan F. Boyce^9^, David L. Boyle^10^, S. Louis Bridges Jr^11^, Jane H. Buckner^12^, Vivian P. Bykerk^7^, Edward DiCarlo^13^, James Dolan^14^, Thomas M. Eisenhaure^4^, Gary S. Firestein^10^, Susan M. Goodman^7^, Ellen M. Gravallese^5^, Peter K. Gregersen^15^, Joel M. Guthridge^16^, Maria Gutierrez-Arcelus^1,2,3,4,5^, Nir Hacohen^4^, V. Michael Holers^17^, Laura B. Hughes^11^, Lionel B. Ivashkiv^18,19^, Eddie A. James^12^, Judith A. James^16^, A. Helena Jonsson^5^, Josh Keegan^14^, Stephen Kelly^20^, Yvonne C. Lee^21^, James A. Lederer^14^, David J. Lieb^4^, Arthur M. Mandelin II^21^, Mandy J. McGeachy^22^, Michael A. McNamara^7^, Nida Meednu^9^, Larry Moreland^22^, Jennifer P. Nguyen^22^, Akiko Noma^4^, Dana E. Orange^23^, Harris Perlman^21^, Costantino Pitzalis^24^, Javier Rangel-Moreno^24^, Deepak A. Rao^5^, Mina ohani-Pichavant^25,26^, Christopher Ritchlin^9^, William H. Robinson^25,26^, Karen Salomon-Escoto^27^, Anupamaa Seshadri^14^, Jennifer Seifert^17^, Danielle Sutherby^4^, Darren Tabechian^9^, Jason D. Turner^28^, Paul J. Utz^25,26^

^9^Division of Allergy, Immunology and Rheumatology, Department of Medicine, University of Rochester Medical Center, Rochester, NY, USA.

^10^Department of Medicine, Division of Rheumatology, Allergy and Immunology, University of California, San Diego, La Jolla, CA, USA.

^11^Division of Clinical Immunology and Rheumatology, Department of Medicine, Translational Research University of Alabama at Birmingham, Birmingham, AL, USA.

^12^Translational Research Program, Benaroya Research Institute at Virginia Mason, Seattle, WA, USA.

^13^Department of Pathology and Laboratory Medicine, Hospital for Special Surgery, New York, NY, USA.

^14^Department of Surgery, Brigham and Women’s Hospital and Harvard Medical School, Feinstein Boston, MA, USA.

^15^Feinstein Institute for Medical Research, Northwell Health, Manhasset, NY, USA.

^16^Department of Arthritis & Clinical Immunology, Oklahoma Medical Research Foundation, Oklahoma City, OK, USA.

^17^Division of Rheumatology, University of Colorado School of Medicine, Aurora, CO, USA.

^18^Graduate Program in Immunology and Microbial Pathogenesis, Weill Cornell Graduate School of Medical Sciences, New York, NY, USA.

^19^David Z. Rosensweig Genomics Research Center, Hospital for Special Surgery, New York, NY, USA.

^20^Department of Rheumatology, Barts Health NHS Trust, London, UK.

^21^Division of Rheumatology, Department of Medicine, Northwestern University Feinberg School of Medicine, Chicago, IL, USA.

^22^Division of Rheumatology and Clinical Immunology, University of Pittsburgh School of Medicine, Pittsburgh, PA, USA.

^23^The Rockefeller University, New York, NY, USA.

^24^Centre for Experimental Medicine & Rheumatology, William Harvey Research Institute, Queen Mary University of London, London, UK.

^25^Division of Immunology and Rheumatology, Department of Medicine, Stanford University School of Medicine, Palo Alto, CA, USA.

^26^Immunity, Transplantation, and Infection, Stanford University School of Medicine, Stanford, CA, USA.

^27^Division of Rheumatology, Department of Medicine, University of Massachusetts Medical School, Worcester, MA, USA

^28^Rheumatology Research Group, Institute for Inflammation and Ageing, NIHR Birmingham Biomedical Research Center and Clinical Research Facility, University of Birmingham, Queen Elizabeth Hospital, Birmingham, UK.

## Statistical analysis

For all the analysis and plots, sample sizes and measures of center and confidence intervals (mean ± SD or SEM), and statistical significance are presented in the figures, figure legends, and in the text. Results were considered statistically significant when *P* < 0.05 by Bonferroni correction and indicated in figure legends and text.

## DATA AVAILABILITY

Upon acceptance, all single-cell sequencing data will be made available on GEO. Upon acceptance, source code to reproduce analyses will be made available on GitHub.

## Reference

1. McInnes, I. B. & Schett, G. The pathogenesis of rheumatoid arthritis. N. Engl. J. Med. 365, 2205–2219 (2011).

2. Neurath, M. F. Cytokines in inflammatory bowel disease. Nat. Rev. Immunol. 14, 329–342 (2014).

3. Liu, J. et al. Overlapping and discrete aspects of the pathology and pathogenesis of the emerging human pathogenic coronaviruses SARS-CoV, MERS-CoV, and 2019-nCoV. J. Med. Virol. 92, 491–494 (2020).

4. Liao, M. et al. Single-cell landscape of bronchoalveolar immune cells in patients with COVID-19. Nat. Med. (2020) doi:10.1038/s41591-020-0901-9.

5. Wen, W. et al. Immune cell profiling of COVID-19 patients in the recovery stage by single-cell sequencing. Cell Discov 6, 31 (2020).

6. Huang, C. et al. Clinical features of patients infected with 2019 novel coronavirus in Wuhan, China. Lancet 395, 497–506 (2020).

7. Lucas, C. et al. Longitudinal analyses reveal immunological misfiring in severe COVID-19. Nature (2020) doi:10.1038/s41586-020-2588-y.

8. He, W., Kapate, N., Shields, C. W., 4th & Mitragotri, S. Drug delivery to macrophages: A review of targeting drugs and drug carriers to macrophages for inflammatory diseases. Adv. Drug Deliv. Rev. (2019) doi:10.1016/j.addr.2019.12.001.

9. Kinne, R. W., Bräuer, R., Stuhlmüller, B., Palombo-Kinne, E. & Burmester, G. R. Macrophages in rheumatoid arthritis. Arthritis Res. 2, 189–202 (2000).

10. Ma, W.-T., Gao, F., Gu, K. & Chen, D.-K. The Role of Monocytes and Macrophages in Autoimmune Diseases: A Comprehensive Review. Front. Immunol. 10, 1140 (2019).

11. Papalexi, E. & Satija, R. Single-cell RNA sequencing to explore immune cell heterogeneity. Nat. Rev. Immunol. 18, 35–45 (2018).

12. Zhang, F. et al. Defining inflammatory cell states in rheumatoid arthritis joint synovial tissues by integrating single-cell transcriptomics and mass cytometry. Nat. Immunol. (2019) doi:10.1038/s41590-019-0378-1.

13. Kuo, D. et al. HBEGF+ macrophages in rheumatoid arthritis induce fibroblast invasiveness. Sci. Transl. Med. 11, (2019).

14. Smillie, C. S. et al. Intra- and Inter-cellular Rewiring of the Human Colon during Ulcerative Colitis. Cell vol. 178 714–730.e22 (2019).

15. Korsunsky, I. et al. Fast, sensitive and accurate integration of single-cell data with Harmony. Nat. Methods (2019) doi:10.1038/s41592-019-0619-0.

16. Stuart, T. & Satija, R. Integrative single-cell analysis. Nat. Rev. Genet. (2019) doi:10.1038/s41576-019-0093-7.

17. Hie, B., Bryson, B. & Berger, B. Efficient integration of heterogeneous single-cell transcriptomes using Scanorama. Nat. Biotechnol. (2019) doi:10.1038/s41587-019-0113-3.

18. Haghverdi, L., Lun, A. T. L., Morgan, M. D. & Marioni, J. C. Batch effects in single-cell RNA-sequencing data are corrected by matching mutual nearest neighbors. Nat. Biotechnol. 36, 421–427 (2018).

19. Butler, A., Hoffman, P., Smibert, P., Papalexi, E. & Satija, R. Integrating single-cell transcriptomic data across different conditions, technologies, and species. Nat. Biotechnol. (2018) doi:10.1038/nbt.4096.

20. Polański, K. et al. BBKNN: fast batch alignment of single cell transcriptomes. Bioinformatics 36, 964–965 (2020).

21. Tran, H. T. N. et al. A benchmark of batch-effect correction methods for single-cell RNA sequencing data. Genome Biol. 21, 12 (2020).

22. Martin, J. C. et al. Single-Cell Analysis of Crohn’s Disease Lesions Identifies a Pathogenic Cellular Module Associated with Resistance to Anti-TNF Therapy. Cell 178, 1493–1508.e20 (2019).

23. Stephenson, W. et al. Single-cell RNA-seq of rheumatoid arthritis synovial tissue using low-cost microfluidic instrumentation. Nat. Commun. 9, 791 (2018).

24. Arazi, A. et al. The immune cell landscape in kidneys of patients with lupus nephritis. Nat. Immunol. 20, 902–914 (2019).

25. Reyfman, P. A. et al. Single-Cell Transcriptomic Analysis of Human Lung Provides Insights into the Pathobiology of Pulmonary Fibrosis. Am. J. Respir. Crit. Care Med. (2018) doi:10.1164/rccm.201712-2410OC.

26. Blondel, V. D., Guillaume, J.-L., Lambiotte, R. & Lefebvre, E. Fast unfolding of communities in large networks. arXiv [physics.soc-ph] (2008).

27. McInnes, L., Healy, J. & Melville, J. UMAP: Uniform Manifold Approximation and Projection for Dimension Reduction. arXiv[stat.ML] (2018).

28. Stoeckius, M. et al. Cell Hashing with barcoded antibodies enables multiplexing and doublet detection for single cell genomics. Genome Biol. 19, 224 (2018).

29. Dallagi, A. et al. The activating effect of IFN-γ on monocytes/macrophages is regulated by the LIF-trophoblast-IL-10 axis via Stat1 inhibition and Stat3 activation. Cell. Mol. Immunol. 12, 326–341 (2015).

30. Fonseka, C. Y. et al. Mixed-effects association of single cells identifies an expanded effector CD4+ T cell subset in rheumatoid arthritis. Sci. Transl. Med. 10, (2018).

31. Luo, W. et al. Targeting JAK-STAT Signaling to Control Cytokine Release Syndrome in COVID-19. Trends Pharmacol. Sci. 41, 531–543 (2020).

32. Wang, B. X. & Fish, E. N. Global virus outbreaks: Interferons as 1st responders. Semin. Immunol. 43, 101300 (2019).

33. Davoudi-Monfared, E. et al. Efficacy and safety of interferon beta-1a in treatment of severe COVID-19: A randomized clinical trial. medRxiv 2020.05.28.20116467 (2020).

34. Nile, S. H. et al. COVID-19: Pathogenesis, cytokine storm and therapeutic potential of interferons. Cytokine Growth Factor Rev. 53, 66–70 (2020).

35. Ye, Q., Wang, B. & Mao, J. The pathogenesis and treatment of the ‘Cytokine Storm’ in COVID-19. J. Infect. 80, 607–613 (2020).

36. Cao, Y. et al. Ruxolitinib in treatment of severe coronavirus disease 2019 (C0VID-19): A multicenter, single-blind, randomized controlled trial. J. Allergy Clin. Immunol. 146, 137–146.e3 (2020).

37. Ahmed, A. et al. Ruxolitinib in adult patients with secondary haemophagocytic lymphohistiocytosis: an open-label, single-centre, pilot trial. Lancet Haematol 6, e630–e637 (2019).

38. Bray, N. L., Pimentel, H., Melsted, P. & Pachter, L. Near-optimal probabilistic RNA-seq quantification. Nat. Biotechnol. 34, 525–527 (2016).

39. Ritchie, M. E. et al. limma powers differential expression analyses for RNA-sequencing and microarray studies. Nucleic Acids Res. 43, e47 (2015).

