## Supplemental figures and materials for "IFN-*γ* and TNF-*α* drive a *CXCL10*+ *CCL2*+ macrophage phenotype expanded in severe COVID-19 and other diseases with tissue inflammation"

#### Methods

##### Integration of scRNA-seq profiles from multiple datasets

*Data collection and aggregation*. To build a multi-tissue immune cell reference, we obtained the raw FASTQ files and raw count matrices from the following publicly available datasets: RA synovial cells from dbGaP (phs001457.v1.p1)[12] and dbGaP (phs001529.v1.p1) [23], SLE kidney cells from dbGaP (phs001457.v1.p1)[24], UC colon cells from Single Cell Portal (SCP259)[14], CD ileum cells from GEO (GSE134809)[22], interstitial and pulmonary lung disease from GEO (GSE122960)[25], and COVID-19 and healthy BALF cells from GEO (GSE145926)[4]. For the FASTQs that we obtained, we used Kallisto [38] to map the raw reads to the same kallisto index generated from GRCh38 Ensembl v100 FASTA files. We pseudo-aligned FASTQ files to this reference, corrected barcodes, sorted BUS files and counted UMIs to generate UMI-count matrices. We aggregated all the cell barcodes from 125 donor samples into one matrix. We performed consistent QC to remove the cells that expressed fewer than 500 genes or with more than 20% of the number of UMIs mapping to the mitochondrial genes, resulting in 307,084 cells in total. The number of donor samples and cells that passed QC for each tissue source, disease status, and technology are shown in **Supplementary Table 1**.

*Normalization, scaling, and feature selection*. We aggregated all samples on the overlapped 17,054 genes. We then normalized each cell to 10,000 reads, and then log-transformed the normalized data. We then selected the top 1,000 most highly variable genes based on dispersion within each dataset. Based on the pooled highly variable genes, we then scaled the aggregated data matrix to have mean 0 and variance 1. We normalized the expression matrix using L2 norm and calculated Cosine distance.

*Dimensionality reduction and batch effect correction*. To minimize the effect of integration from multiple datasets with different contributing cell numbers, we use weighted principal component analysis (PCA) and used top 20 PCs for follow-up analysis. The summation of the weights is equal for each separate tissue source. Then we corrected the batch effects from three levels: sequencing technology, tissue source, and donor sample simultaneously using weighted Harmony [15]. We use default parameters and also specified theta = 2 for three variables including tissue source, technology and sample, respectively, max.iter.cluster = 30, and max.iter.harmony = 20. In the weighted Harmony, we use the same weights from the weighted PCA by setting more weights for the cells from a tissue source with fewer number of cells, and less weights for the cells from a tissue source with a greater number of cells. As outputs, we obtained batch corrected PC embeddings where the effects from different single-cell datasets and donors are removed in the low-dimensional PC space.

*Quantitative evaluation*.

Variance explained from different sources: To quantitatively measure the mixture of batch effects after correction, we estimated the sources of variance explained in the gene expression from original pre-defined immune cell type, tissue origin, technology, and donor sample in the first ten principal component embeddings. We used the R package limma to fit the model and anova to compute the variance explained percentage:

$$principal\ component\ \sim celltype\ +\ tissue\ +\ technology\ +\ sample.$$

LISI score: In the meanwhile, we used a LISI (local inverse Simpson's Index) metric to measure the mixture levels of batch labels based on local neighbors chosen on a specific perplexity [15,21] . Specifically, we build Gaussian distribution of neighborhoods and compute these local distributions of batch probabilities $p(b)$ using perplexity 30 on the top 20 principle components. Then we calculated the inverse Simpson's Index:

$$1/\sum_{b=1}^{B} p(b),$$

where the LISI score ranges from 1 (unmixed of batches) to B (the maximum score is the total number of levels in the categorical batch variable). Here batch can be tissue source, donor sample, and sequencing technology.

*Graph-based clustering*. We then applied unbiased graph-based clustering (Louvain [26]) on the top 20 batch corrected PCs at various resolution levels (0.2, 0.4, 0.6, 0.8, 1.0). We use 0.4 as the resolution value to gain the biological interpretations that make most sense. Then, we furthermore performed dimensionality reduction using UMAP [27] on the top 20 batch corrected PC embeddings.

*Single-cell Differential expression analysis*. We identified cell type cluster marker genes by comparing cells within one cluster with all the other clusters. We calculated several criteria to decide statistically significant marker genes: 1) Fold-change, 2) AUC, 3) Wilcoxon rank-sum test and Bonferroni corrected *P*, and 4) percent of cells were expressed within a specific cluster. We tested all the genes that were detected at more than 100 cells with non-zero UMI counts.

*Identification of major immune cell type clusters.* We carefully annotated each identified cluster using two ways. First, we mapped the original publication labels [12,14,22] to our UMAP embeddings to show we are able to reproduce original cell type labels (**Supplementary Figure**

**1**). Second, we annotated the identified clusters using cell type lineage marker genes, for example, T cells (*CD3D*), NK cells (*NKG7*), B cells (*MS4A1*), plasma cells (*MZB1*), macrophages (*FCGR3A* and *CD14*), dendritic cells (DCs, *CD1C*), mast cells (*TPSAB1*), and cycling cells (*MKI67*).

**Cell culture for human blood–derived macrophages and synovial fibroblasts**

We obtained human leukocyte enriched whole blood samples from 4 healthy blood donors from New York Blood Center and purified peripheral blood mononuclear cells (PBMC) from each using Ficoll gradient centrifugation. We isolated CD14+ monocytes from each sample using human CD14 microbeads (Miltenyi Biotec) and differentiated into blood-derived macrophages for 1 day at 37° C in macrophage CSF (M-CSF; 10 ng/ml) (PeproTech) and RPMI 1640 medium (Corning) supplemented with 10% defined fetal bovine serum (FBS) (HyClone), 1% Penicillin-Streptomycin (Thermo Fisher Scientific), and 1% L-glutamine (Thermo Fisher Scientific) in a 6-well plate a concentration of 1.2 million cells/mL.

In parallel, we obtained human synovial fibroblasts derived from deidentified synovial tissues of RA patients undergoing arthroplasty (HSS IRB no. 14-033). We cultured fibroblasts in alpha minimum essential medium (a-MEM) (Gibco) supplemented with 10% premium FBS (R&D Systems Inc), 1% Penicillin-Streptomycin (Thermo Fisher Scientific), and 1% L-glutamine (Thermo Fisher Scientific) for 4 to 6 passages. To create each trans-well, we seeded the mesh of polyester chambers with 0.4- um pores (Corning) with either 200,000 synovial fibroblasts or without fibroblasts for 1 day at 37° C.

The following day, we suspended each trans-well - 3 with fibroblasts and 6 without fibroblasts per donor - above one well of cultured macrophages. Those with fibroblasts had a fibroblast--to--macrophage ratio of 1:15. In total, we created 9 wells per donor. Next, we added either IFN-$\beta$

(200 pg/mL), IL-4 (20 ng/ mL), TNF-$\alpha$ (20 ng/mL), and/ or IFN-$\gamma$ (5 ng/mL) to each trans-well and underlying plate per donor. All plates were incubated at 37° C for 19 hours.

**RNA library preparation and sequencing**

Next, we applied a modified version of the staining protocol from CITE-seq, using only Totalseq$^{TM}$-A Hashing antibodies from Biolegend. We harvested macrophages from each well and aliquoted one fifth of the cells, ~750,000 cells per condition, for staining in subsequent steps. We washed the cells in filtered labeling buffer (PBS with 1% BSA) and resuspended in 50 ul of labeling buffer with Human TruStain FcX$^{TM}$ (Biolegend Cat #422302, 5 ul per stain) for 10 min at 4° C. Next, we added 50 ul of labeling buffer with 1.6 ng/uL of a Total-seq hashtag (1, 2, 4, 5, 6, 7, 8, 9, or 12) per condition per donor for 25 minutes at 4° C. Complete Hashtag details are listed in **Supplementary Figure 7a** and **Supplementary Table 4**. Next, we washed all samples in 2 mL, 1 mL, and 1 mL of labeling buffer, sequentially. We counted the remaining cells using a cellometer (Nexcelom Cellometer Auto 1000) and aliquoted the equivalent of 60,000 cells from each condition into one Eppendorf tube per donor. From here we filtered through a 40um mesh and resuspended in PBS with 0.04% BSA to a concentration of 643.7 cells/ ul. We followed the Chromium Single Cell 3' v3 kit (10x Genomics) processing instructions and super-loaded 30,000 cells per lane. We used one lane per donor, with 9 conditions multiplexed per donor sample. After cDNA generation, samples were shipped to the Brigham and Women’s Hospital Single Cell Genomics Core for cDNA amplification and sequencing. Pairs of libraries were pooled and sequenced per lane on an Illumina NovaSeq S2 with paired-end 150 base-pair reads (**Supplementary Table 5**).

**Processing FASTQ reads into gene expression matrices and cell hashing**

We quantified mRNA and antibody unique molecular identifiers (UMI) counts, respectively. Cellranger v3.1.0 was used to process the raw BCL files and produce a final gene by cell barcode

UMI count matrix. First, raw BCL files were demultiplexed using cellranger mkfastq to generate FASTQ files with default parameters. Then, these FASTQ files were demultiplexed using Cellranger v3.1.0 and aligned to the GRCh38 human reference genome. Gene/antibody reads were quantified simultaneously using cellranger count. Cell barcodes and UMIs were extracted for gene/hashtag antibodies for each run.

For quality control of the cells, we first performed mRNA-level cell QC and then hashtag-level QC. For the mRNA-level QC, we removed the cells that expressed fewer than 1,000 genes or more than 10% of the number of UMIs mapping to the mitochondrial genes. For the hashtag QC, we removed the cells whose proportion of UMIs for the most abundant hashing antibody is less than 90%; and removed the cells whose ratio of the second most-abundant and first most-abundant antibody is greater than 10%. After filtering, each cell was assigned a hashing antibody and donor sample on the most abundant hashing antibody barcode. After QC, we obtained 9,399 cells, 8,775 cells, 4,622, and 3,027 cells for each of the 4 donor samples. Then we normalized each cell based on the total number of UMIs and log-transformed it.

**Pseudo-bulk differential expression analysis**

To identify the robust cluster marker genes that are shared between diseases for each identified macrophage and monocyte-specific cluster, we performed pseudo-bulk analysis by summing the raw UMI counts for each gene across cells from the same donor sample, tissue source, and cluster assignment. Then we log normalized counts for each gene in each pseudo-bulk sample into counts per million (CPM). We removed the tissue-specific batch effect by setting tissue source as covariate using R function removeBatchEffect from limma [39] package. Then we identified the cluster (*CXCL10*+ *CCL2*+ and *FCN1*+ states) marker genes using Wilcoxon Rank Sum Test and AUC on the pseudo-bulk tissue batch corrected matrix. We consider genes with

AUC greater than 0.6, and *P* smaller than the Bonferroni correction threshold $10^{-5}$ (0.05/5,000 tested highly variable genes) are statistically significant.

**Linear modeling for experimental stimulation-specific genes from cell culture single-cell profiles**

We clustered single-cell profiles from human blood-derived macrophage hashtag experiments and identified stimulation-driven clusters. To more accurately identify the gene signatures that are specific to each of the eight stimulation, we use linear models to test each gene for differential normalized gene expression across contrasts of interest. Specifically, we fit the following models:

$$gene\_expression \sim stim + 1|sample + nUMI,$$

where $stim$ is a categorical variable that represents eight stimuli and an untreated status, $1|sample$ is the random effect of the 4 replicated donor samples, $nUMI$ (unique molecular identifier) represents the technical cell-level fixed effect. We obtained the fold-change, *T* and *P* value, and Bonferroni corrected *P* as measurements for each tested gene signature for each applied condition. We then generated a list of differentially expressed genes whose fold-change is greater than 2 and *P* is smaller than the Bonferroni correction threshold $10^{-7}$ (0.05/7,000 highly variable genes x 9 conditions) for each stimulus condition.

**Testing integrative macrophage clusters for association with severe/inflamed status**

We tested the association of each macrophage cluster with severe/inflamed status compared to healthy with MASC (mixed-effects modeling of associations of single cells) [30]. We fit a logistic regression model for each identified cluster within one tissue and set the nUMIs and percent of MT content (% MT) as cell-level fixed effect, and donor sample as random effect covariates:

$$log[\frac{Y_{i,c}}{1-Y_{i,c}}] = \beta_{case}X_{i,case} + \beta_{tech1}X_{i,tech1} + \beta_{tech2}X_{i,tech2} + (\varphi_d | \, d \,),$$

where $Y_{i,j}$ is the odds of cell $i$ in cluster $c$, $\beta_{case}$ is the effect log(Odds ratio) for case (Severe COVID-19)-control (healthy) status, $\beta_{tech1}$ is a vector of technical cell-level (nUMIs) covariate, $\beta_{tech2}$ is a vector of technical cell-level (% mitochondrial genes) covariate, $X_i$ is the values for cell $i$ in technology as appropriate, and $(\varphi_d | d)$ is the random effect of donor $d$. Thus, we used this logistic regression model to test for differentially abundant macrophage clusters associated with severe COVID-19 by correcting for the technical cell-level and donor-level covariates. Similarly, we also tested for differentially abundant macrophage clusters associated with inflamed CD compared to non-inflamed CD, RA compared to OA, inflamed UC compared to healthy colon, respectively accounting for technical cell-level and donor-level covariates. We generated log likelihood-ratio test MASC *P* and odds ratio for each tested cluster and used Bonferroni correction to report the macrophage clusters that are statistically significantly abundant in severe/inflamed samples compared to healthy or non-inflamed controls.

#### Supplementary Figure Legends

**Supplementary Figure 1.** Overall integration of immune cells from multiple scRNA-seq datasets. **a**. We describe the diversity of the single-cell sequencing technologies using nGene (number of genes detected) and nUMI (number of unique molecular identifiers). **b**. In the PC1 and PC2 and UMAP1 and UMAP2 after batch effect correction, we show cells from different tissue sources separately and color each cell based on disease tissue categories. The same cell types from different tissue sources cluster together in the PCA and UMAP space, respectively. **c**. We are also able to reconstruct the high-resolution of the immune cell subsets in the overall integrative embeddings. We color each cell by the original published cell subset annotations from UC colon and RA synovium [12,14]. **d**. Percent of variance explained for the top 10 PCs are shown.

**Supplementary Figure 2.** Quantify the diversity of different batches before and after batch effect correction. **a**. We quantified the mixture level of donor samples and tissue sources using LISI score for before and after harmony batch effect correction. The LISI scores that measure mixture levels of donor and tissue are increased after using harmony batch effect correction compared to before correction. **b**. We show the disease category, sequencing technology, and tissue source for each cell in the UMAP space before correcting batch effects. Most of the cell clusters are driven by separate tissue sources rather than shared cell type populations before batch correction.

**Supplementary Figure 3.** Integrative analysis of macrophages from multiple scRNA-seq datasets from human tissue. **a**. We show the gene expression of macrophage cluster marker genes in the UMAP space. **b**. Original macrophage subset annotations [12,14] are projected into the integrative UMAP embeddings. The previously identified inflammatory macrophages from UC colon and RA synovium are colored and labeled. Cells from RA (red) and OA (black) synovium are colored. **c**. Individual samples from healthy, mild and severe COVID-19 are shown separately based on the same integrative UMAP coordinates. **d**. We use LISI score to quantify the mixture level of tissue source and donor sample, and original macrophage subset identification before and after batch correction.

**Supplementary Figure 4.** Heterogeneity of shared inflammatory macrophages from multiple tissues. **a**. Proportion of expressing (non-zero) inflammatory cytokines and genes from inflammatory macrophages in severe COVID-19 are higher compared to healthy BALF. Genes that are highly expressed in the *CXCL10+ CCL2+* inflammatory macrophages are highlighted in orange. **b**. Distribution of macrophages from each disease tissue along the identified PC1, which was identified by performing PCA analysis on the inflammatory macrophages. PC1 reflects a gradient from *FCN1+* to *CXCL10+ CCL2+*. **c**. Heterogeneity of the identified *CXCL10+ CCL2+* inflammatory macrophages correlates with *IL1B* expression. **d**. We show the expression of gene *FCN1*, *CXCL10, IL1B,* and *CCL2* for different disease tissue sources.

**Supplementary Figure 5.** Differential expression analysis comparing inflammatory macrophages with non-inflammatory macrophages within each individual tissue source at the single-cell level. Log-transformed fold change and *P* for each gene are shown. Within RA synovium, we conducted differential gene expression analysis by comparing *FCN1+* macrophages with *MRC1+ FABP4+* macrophages from the same RA synovium, and also *CXCL10+ CCL2+* macrophages with *MRC1+ FABP4+* macrophages from the same RA synovium. We performed similar differential gene expression analysis for UC colon and COVID-19 BALF.

**Supplementary Figure 6**. Bulk RNA-seq gene expression of inflammatory macrophage-associated genes using CD45+ CD14+ flow sorted macrophages from RA and OA synovium (ImmPort SDY998). We show the expression of genes that are specific to the *CXCL10+ CCL2+* and *FCN1+* states from each bulk RNA-seq sample in the leukocyte-rich (n=23), leukocyte-poor (n=11), and OA (n=13) groups. TPM denotes transcript count per million. Dots represent samples, lines represent means. Wilcoxon rank-sum test is applied.

**Supplementary Figure 7.** Experimental design and quality control of human blood-derived macrophages stimulated by different conditions. **a**. Schematic plot of experimental design for the human blood-derived macrophages. Seeding macrophages and fibroblasts, addition of cytokines, and single-cell hashing tag steps were included for the experiment. **b**. We filtered cells expressing fewer than 1,000 nGene or with more than 10% of UMIs mapping to mitochondrial genes. We also filtered cells whose proportion of UMIs for the most abundant hashing antibody was less than 90%; and removed cells whose ratio of the second most-abundant to first most-abundant antibody was greater than 10%. **c**. Number of cells that passed stringent QC for each sample. **d**. Heatmap illustrating Z-score of the normalized gene expression for condition-specific genes.

**Supplementary Figure 8.** Integration of tissue-level macrophages and human blood-derived macrophages. **a**. We display macrophages from each tissue source that contribute to each integrative cluster, respectively. Cluster assignments are labeled and colored. **b**. For the blood-derived macrophages, we label each cell using the applied stimulatory condition. We also show the percent and number of stimulated blood-derived macrophages for each stimulatory condition and each cluster. **c**. Heatmap illustrating Z-score of the normalized gene expression of cluster-specific marker genes. **d**. Number of macrophages for each individual sample from healthy, mild and severe COVID-19 BALF groups; and number of macrophages from each stimulatory condition for the blood-derived macrophages. **e**. Gene expression of *CXCL11*, *CXCL10*, *GBP1*, *STAT1*, *TNFSF10*, *IRF1*, *CCL3*, *CCL4*, *CCL2*, and *IDO1* in BALF and blood-derived macrophages in violin plots. Mean of normalized gene expression is marked by a line and each disease status or stimulatory condition by individual coloring.

### Supplementary Figure 1

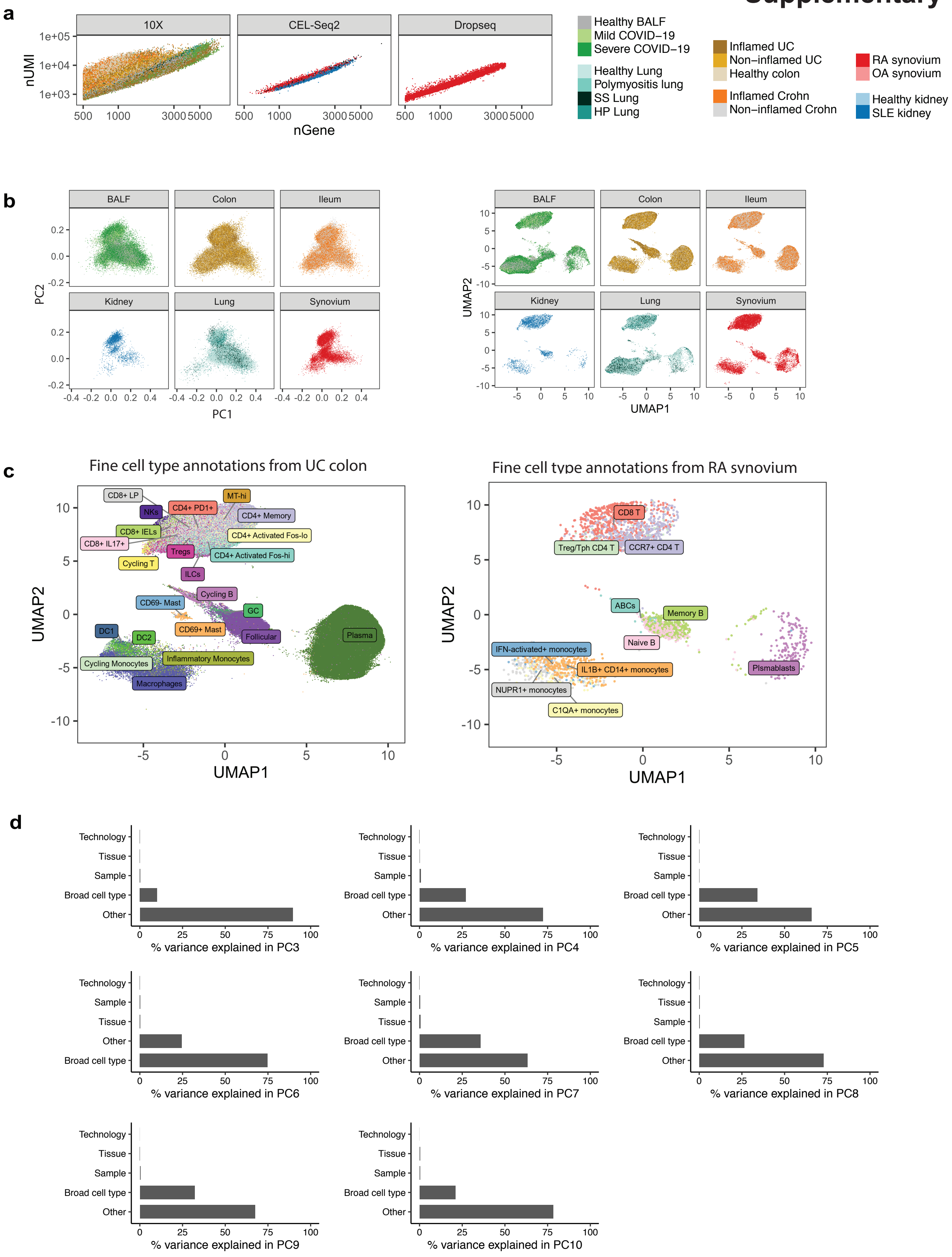

### Supplementary Figure 2

**a**

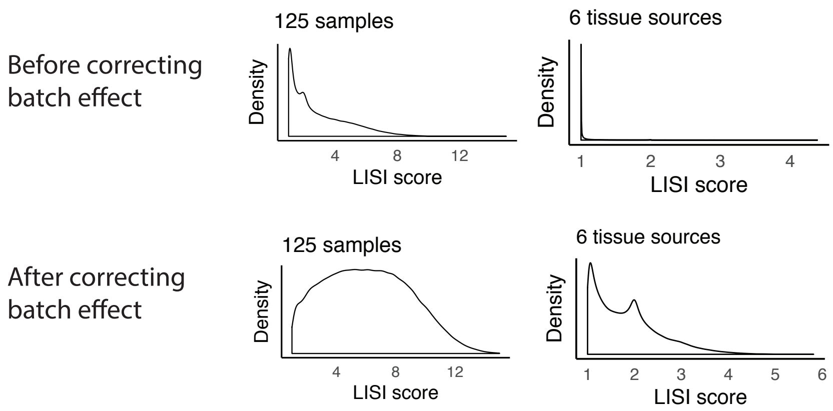


**b**

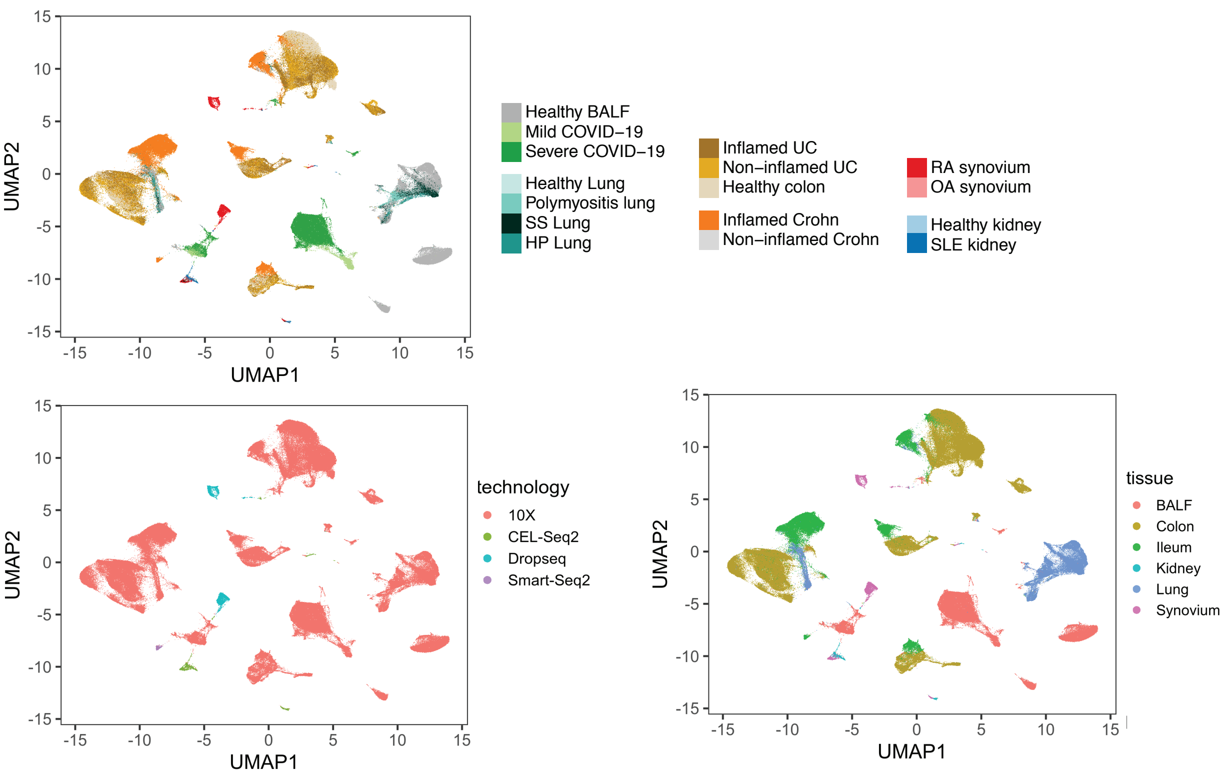

### Supplementary Figure 3

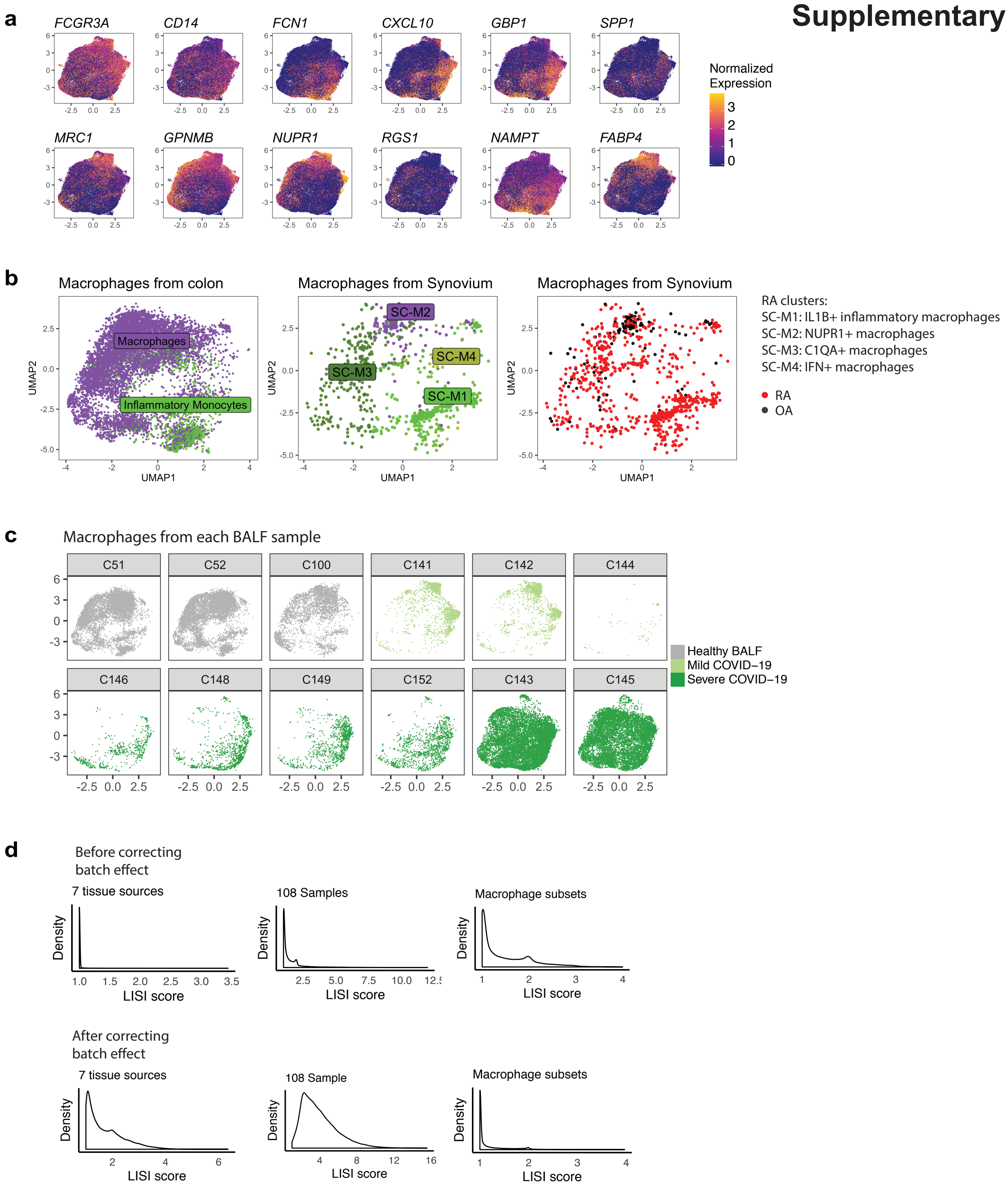

### Supplementary Figure 4

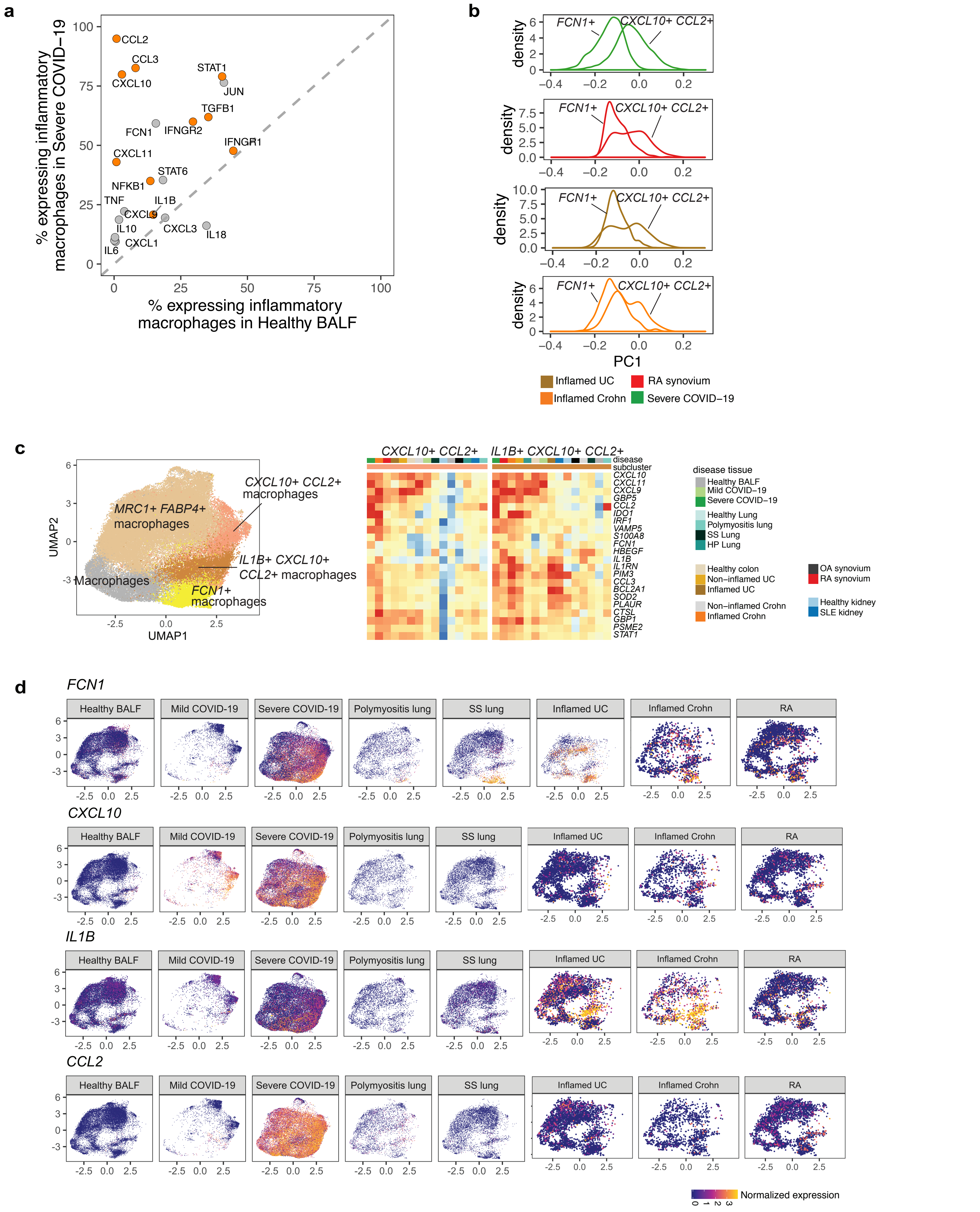

### Supplementary Figure 5

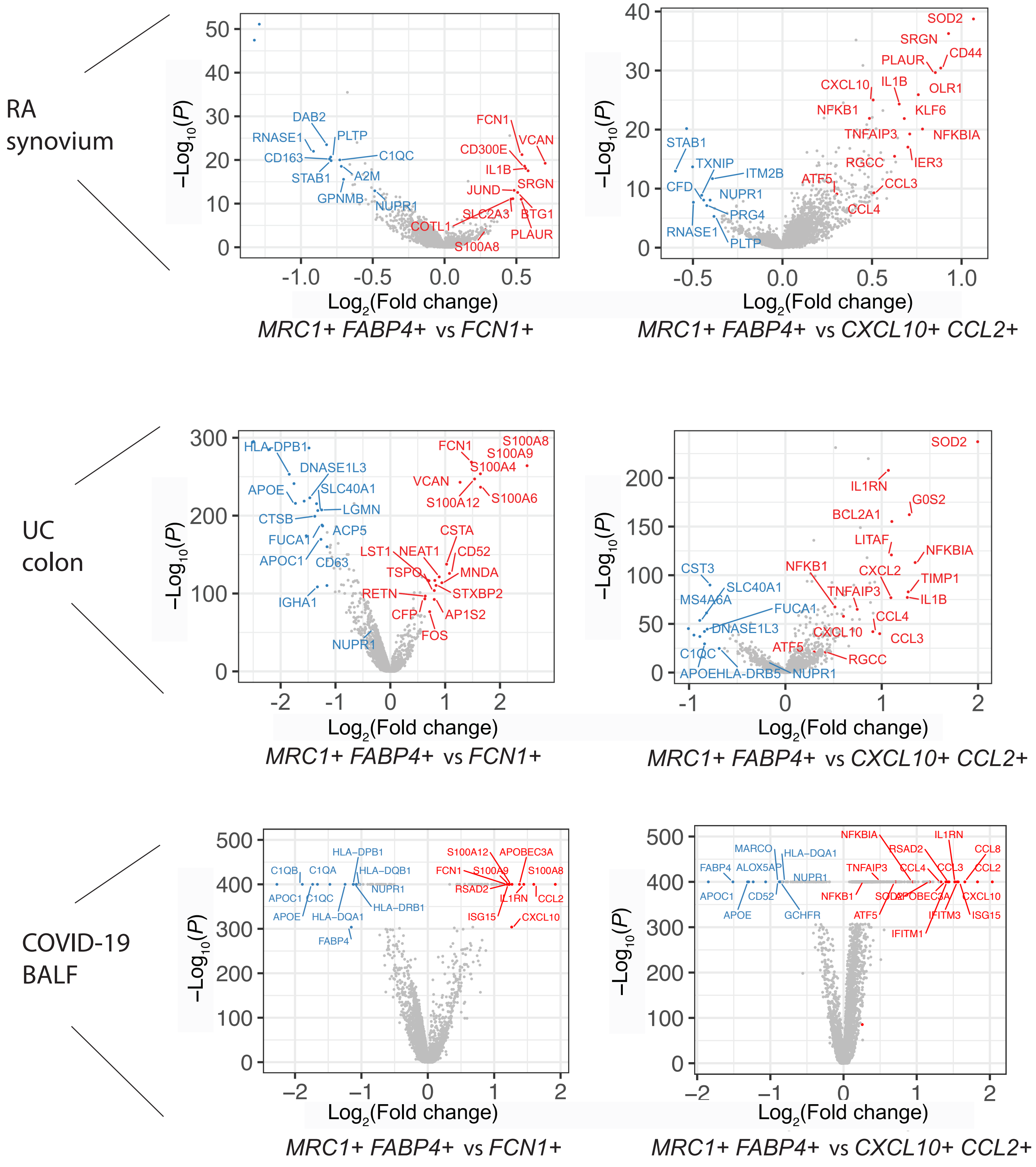

### Supplementary Figure 6

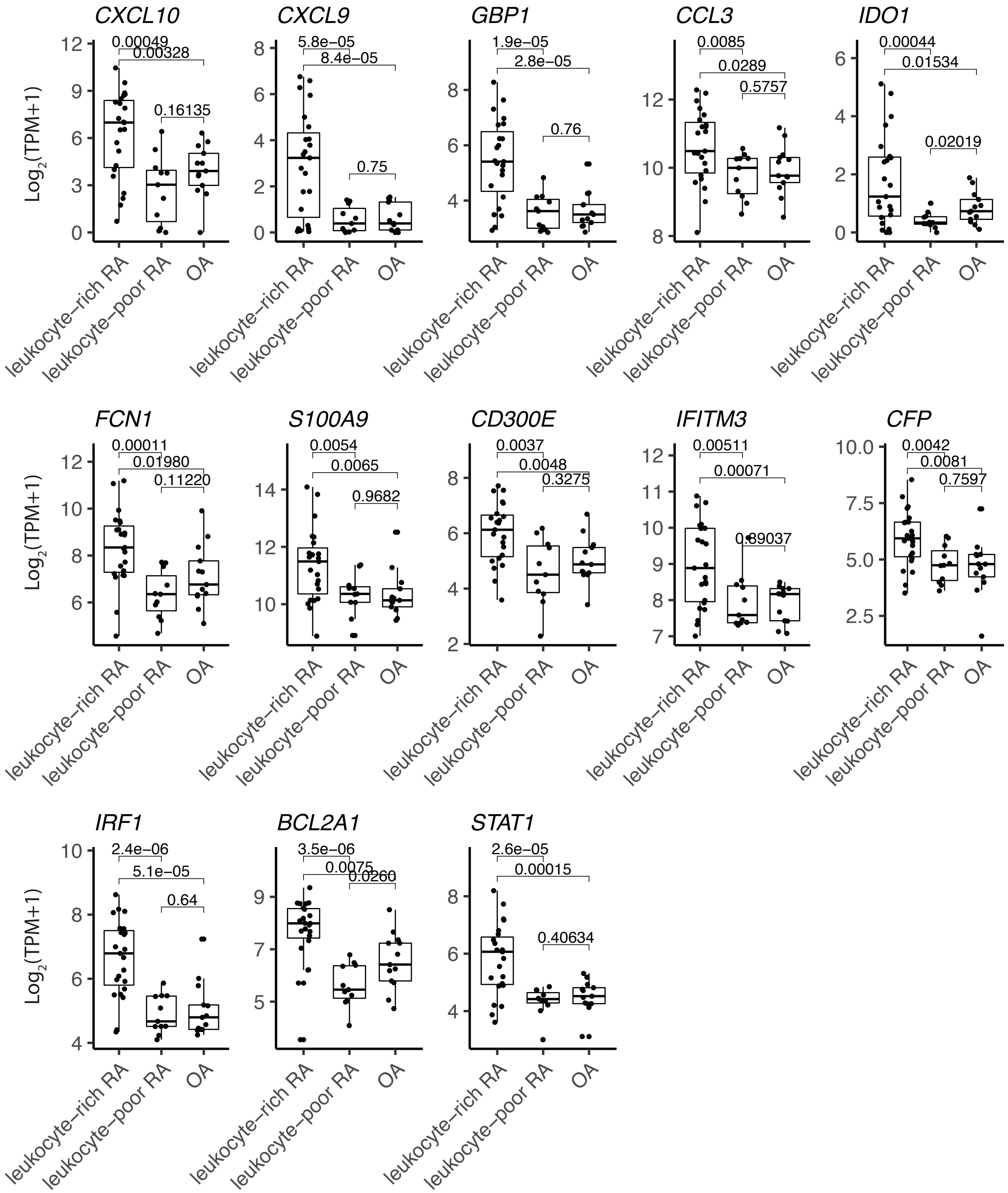

### Supplementary Figure 7

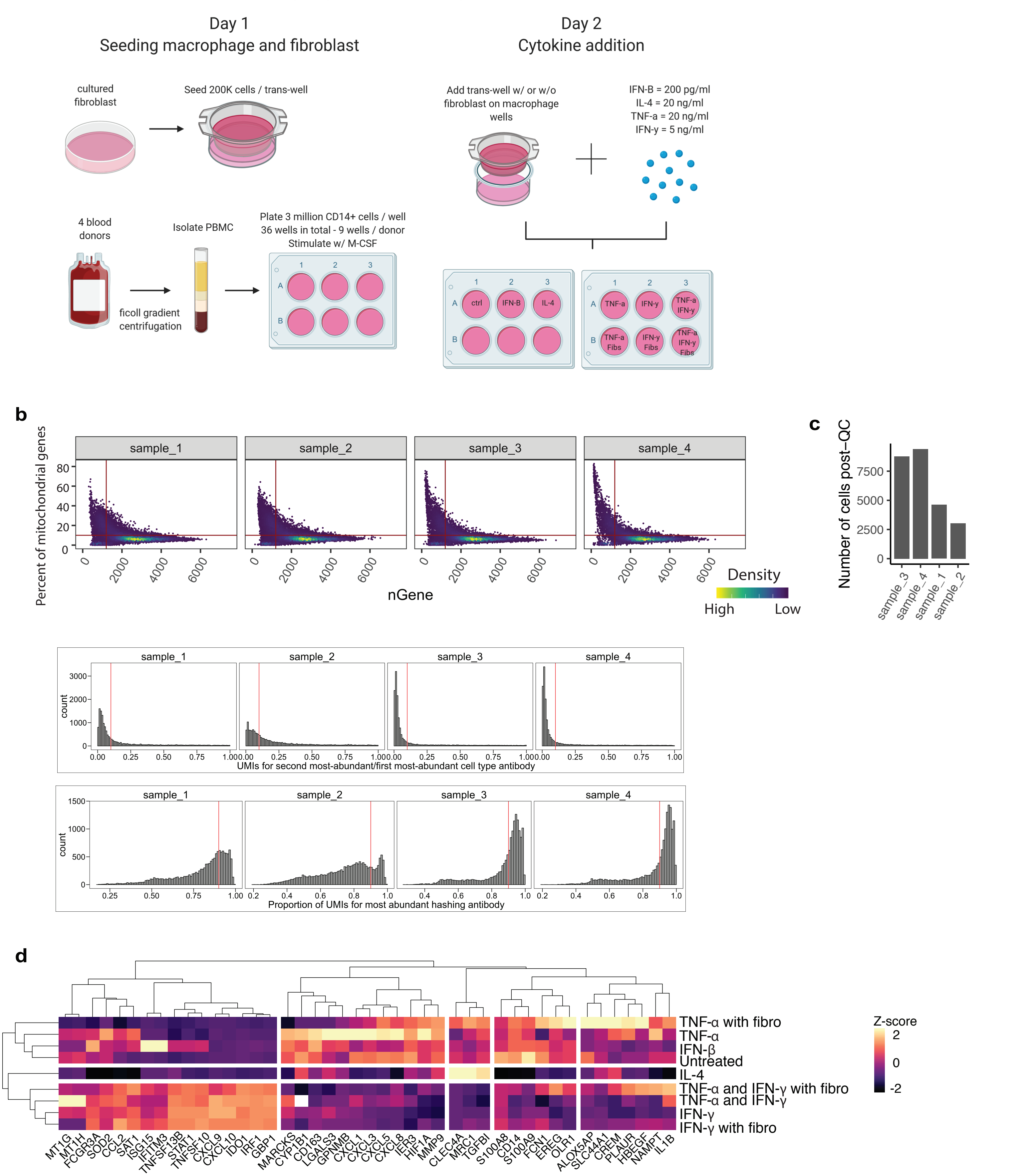

### Supplementary Figure 8

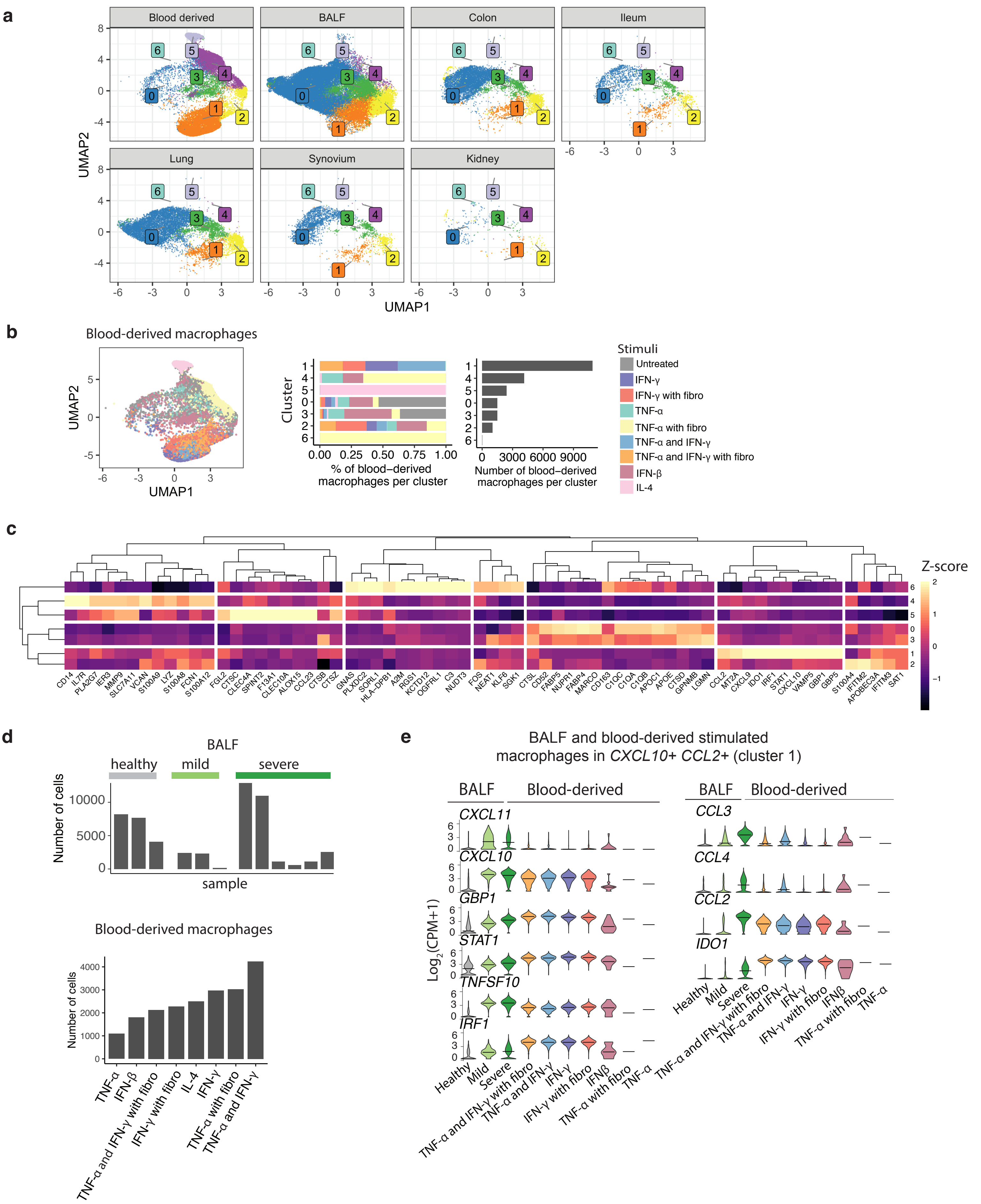
